# MOSAIC: a longitudinal phenotypic clock to dissect organismal aging trajectories in *C. elegans*

**DOI:** 10.64898/2026.04.23.720365

**Authors:** Alexandre Pierre Vaudano, Marie Pierron, Lazar Stojkovíc, Mathieu Membrez, Morgane Bourgeois, Christopher Neal, Myriam Chimen, Lina Verbakel, Matteo Cornaglia, Florence Solari, Laurent Mouchiroud

## Abstract

Interventions that extend lifespan do not necessarily preserve healthspan, the portion of life spent in good health. This disconnect has intensified interest in biological aging clocks as quantitative proxies of organismal health. However, most existing clocks rely on invasive or endpoint measurements, providing static estimates that capture biological age at a single time point and offer limited insight into aging trajectories – the dynamic patterns through which physiological resilience and functional capacity change within individuals over time.

Here we combine standardized, high-frequency imaging of individual *Caenorhabditis elegans* across the lifespan with machine learning to develop MOSAIC (Modular Organismal Signature of Aging In *C. elegans*), a non-invasive phenotypic clock that estimates biological age longitudinally at single-organism resolution. Leveraging ∼3’750 animals, ∼230’000 observations and 29 phenotypic features, MOSAIC predicts biological age with high accuracy and resolves organism-wide aging trajectories at high temporal resolution. Beyond age prediction, MOSAIC decomposes biological age into contributions from distinct physiological modules, enabling mechanistic interpretation of organismal decline.

Applying MOSAIC to natural lifespan variation, dietary restriction, longevity mutants and pharmacological interventions reveals that lifespan extension can emerge through distinct, time-dependent phenotypic trajectories rather than a uniform slowing of aging. Interventions with similar effects on longevity produce divergent biological-age trajectories and distinct combinations of younger and older traits, highlighting context-dependent physiological trade-offs. MOSAIC provides a scalable, non-invasive framework to repeatedly quantify biological age across the lifespan and to compare interventions based on how they reshape aging trajectories.

## Introduction

With populations aging worldwide, identifying genetic, dietary and pharmacological interventions that extend not only lifespan but also healthspan has become a major biomedical priority^1^. Numerous lifespan-extending interventions have been reported across species, from *Caenorhabditis elegans* to humans^2–6^. However, longer life does not necessarily mean healthier life. In *C. elegans*, several long-lived mutants exhibit extended periods of frailty, and the relationship between lifespan and healthspan varies across interventions and genetic backgrounds ^7,8^. In humans, gains in life expectancy have likewise outpaced improvements in healthy life expectancy^9,10^. Together, these findings highlight a key limitation of lifespan as a sole metric of aging and underscore the need to understand how interventions reshape aging trajectories – the dynamic progression of physiological decline within individuals.

Tracking healthspan requires quantifying biological age, a measure of functional state that captures organismal condition more accurately than chronological time. Over the past decade, machine-learning-based aging clocks have emerged as powerful tools to estimate biological age from molecular, clinical and phenotypic data^11–15^, often predicting morbidity, functional decline and mortality risk better than chronological age^16^. Yet most existing clocks rely on cross-sectional or endpoint measurements collected in cohorts, capturing population-level variation rather than the dynamics of aging within individuals^17,18^. As a result, these approaches provide limited insight into whether interventions slow the pace of aging, alter its trajectory, or both – key distinctions for understanding mechanisms that promote healthspan.

The nematode *C. elegans*, with its short lifespan (∼3 weeks) and optical transparency, provides a tractable system for longitudinal, non-invasive measurement of aging. Several molecular aging clocks based on transcriptomic data have achieved accurate prediction of biological age in this organism^19–21^, but require destructive sampling and therefore cannot track individuals continuously across life. Phenotypic clocks based on functional or morphological traits offer a complementary strategy for longitudinal assessment^22–26^, but require large standardized datasets that are difficult to generate reproducibly across laboratories and experimental conditions, as highlighted by initiatives such as the Caenorhabditis Intervention Testing Program²⁷˒²⁸. Here we combine standardized high-frequency imaging with machine learning to develop MOSAIC (Modular Organismal Signature of Aging in *C. elegans*), a longitudinal phenotypic clock that predicts biological age at the level of individuals with high accuracy (median R² = 0.84) and resolves organism-wide aging trajectories from ∼230’000 observations across 29 phenotypic features in ∼3’750 animals. Applying MOSAIC across genetic, dietary and pharmacological interventions reveals that comparable lifespan extensions can arise from distinct, nonlinear biological age trajectories, reflecting independent dynamics of physiological modules. Our work establishes a non-invasive framework to quantify aging trajectories across the lifespan and reveals unexpected diversity in organismal aging dynamics.

## Results

### Automatic worm monitoring with a microfluidics-based device

To systematically profile phenotypic trajectories across the lifespan of *C. elegans*, we used SydLab™ One, a microfluidics-based platform integrating custom-designed chips, automated fluid handling, temperature control, and scheduled imaging at 6x magnification under tightly standardized conditions (Fig. S1a). Each chip comprises 128 microchambers delimited by PDMS pillar filters (Fig. 1a,b). At the start of each experiment, synchronized L4-stage worms were introduced into the microchannels using an optimized loading procedure that confined one to four individuals per chamber. During loading, controlled liquid flow transiently deforms the PDMS pillars, enabling uniform distribution of worms across chambers; upon cessation of flow, the pillars return to their initial spacing, effectively trapping the animals. To maintain stable environmental conditions and prevent progeny accumulation, a bacterial suspension consisting of freeze-dried bacteria was delivered automatically every hour, while newly hatched L1 larvae were flushed out of the device. Images were acquired every 6 hours at a fixed interval (30s) following feeding, ensuring consistent physiological context for longitudinal phenotypic measurement.

**Fig. 1.**
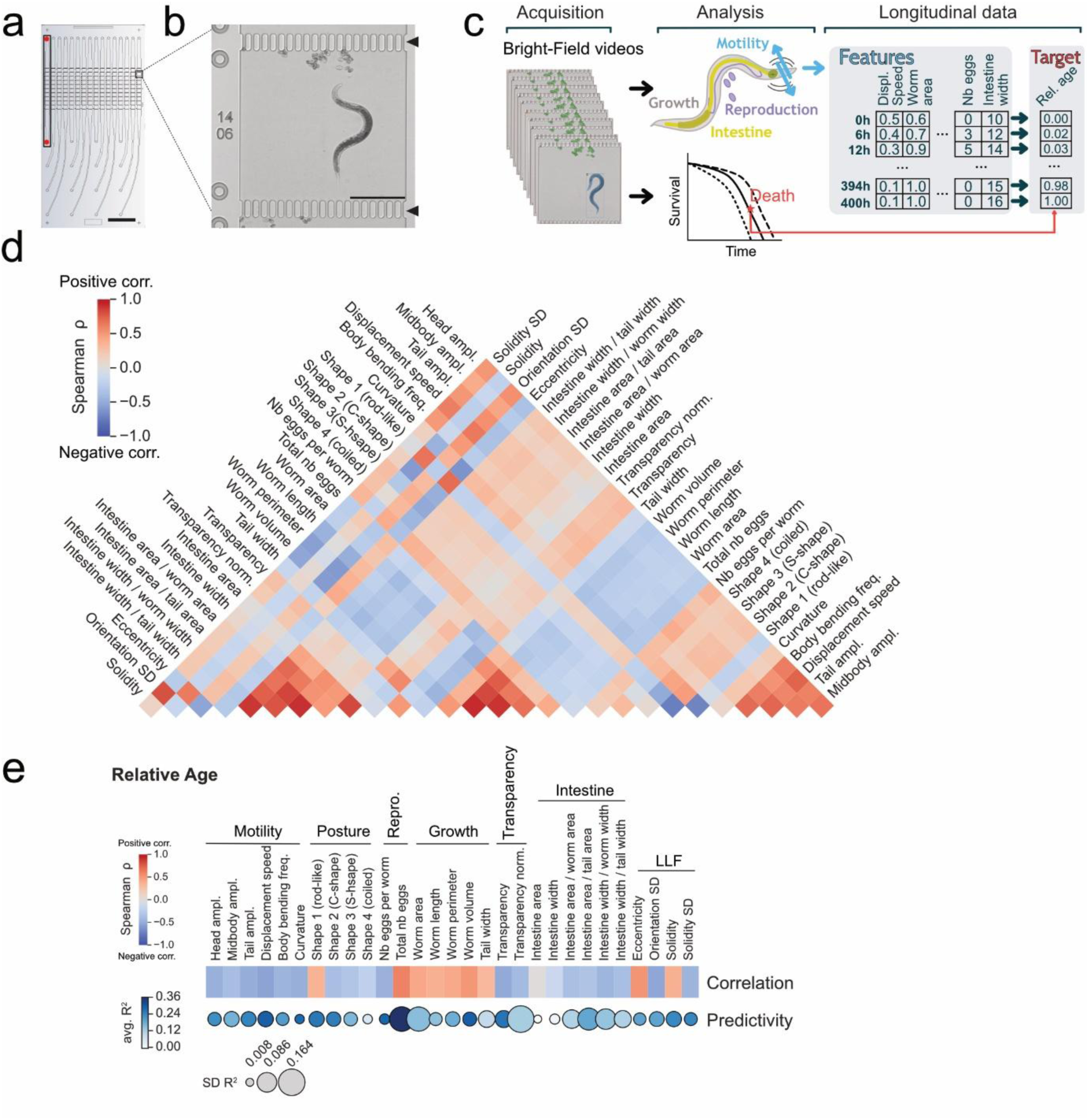
High-temporal resolution analysis of *C. elegans* physiological changes during aging. **a**, Schematic of the device’s microfluidic chip. A chip is composed of 16 independent channels (the black rectangle highlights the first channel) each connected to an inlet and an outlet (red dots). Each channel comprises 8 experimental chambers imaged every 6 hours. Scale bar, 1 cm. **b**, Representative image of an N2 adult worm housed inside a microfluidic chamber taken by the platform (6x Brightfield image). The chamber is delimited by filters made of PDMS pillars (black arrowheads). Scale bar = 500 μM. **c**, schematic explanation of dataset construction, Displ.: displacement. Rel.: relative. **d**, Heatmap of features-to-features Spearman correlation, including data from all ages. **e**, Heatmap of the Spearman correlation of features with relative age and predictivity (R^2^) of relative age for each feature by fitting spline models. avg.: average, SD: standard deviation.

### High-temporal resolution analysis of *C. elegans* physiological changes during aging

Using this microfluidics-based system, we generated an initial longitudinal dataset comprising 2s bright-field (BF) videos acquired every 6 hours from 366 individually housed wild-type (WT) N2 worms, followed from the late L4 stage until death (Fig. S1b, Fig. S2). These videos were analyzed using deep-learning (DL) algorithms to segment worms, eggs, and larvae, from which we extracted 29 quantitative phenotypic features (Fig. 1c, Fig. S1c-e; see methods) that differ in their rate and direction of changes over time. Among them, motility and postural dynamics declined progressively throughout adulthood, reflecting reduced activity and a shift toward straighter body postures, with worms transitioning from active C- and S-shaped postures (shape 2 and 3) to an inactive rod-like shape (shape 1) (Fig. S3a-b). Reproduction followed the expected biphasic pattern, with egg output peaking around day 3 of adulthood before declining sharply, leading to most eggs being laid before day 7 (Fig. S3c). Body size increased during early adulthood and reached a plateau around day 8 (Fig. S3d). Optical transparency exhibited a biphasic trajectory, with rapid early darkening followed by a slow continuous decline (Fig. S3e), likely representing fat accumulation ^27,28^. Intestinal traits displayed multiphasic dynamics. Absolute intestinal size increased until day 5 in parallel with the overall worm growth, then declined; (Fig. S3f) while size-normalized intestinal metrics decreased steadily with age, consistent with progressive intestinal atrophy (Fig. S3g). Finally, low-level features (LLFs) followed similar kinetics to either motility (orientation SD, solidity SD) or posture (eccentricity, solidity) (Fig. S3h). Feature-to-feature correlation across lifespan revealed distinct clusters of positively and negatively associated traits, (Fig. 1d) which identified seven physiological traits (motility, posture, reproduction, growth, transparency, intestine and LLFs) monitored by our pipeline.

We next asked whether age-dependent changes in extracted features could serve as proxies for organismal health and thereby predict life expectancy at any point across the lifespan. We therefore computed the relative age of each worm at each observation time, defined as the ratio of chronological age to individual lifespan (time of death) (see Methods). At the population level, most features were significantly correlated with relative age (Fig. 1e; mean absolute Spearman’s rho = 0.40 ± 0.12 SD), indicating that many phenotypes report progression through the lifespan.

We then assessed the predictive value of each feature separately by fitting spline regression models to estimate the relative age of a worm from single observations (see Methods). Total number of eggs was the most informative single feature predictor (mean R² = 0.36 ± 0.14 SD in 5-fold cross-validation), followed by displacement speed (0.30 ± 0.05) and worm volume (0.30 ± 0.04) (Fig. 1e). These results indicate that, although many traits vary with relative age at the population level, no single phenotype captures individuals’ relative age with high accuracy.

### Different machine learning architectures accurately quantify relative age across the lifespan

To capture organismal state in a multidimensional manner, we combined multiple phenotypic features. We first applied principal component analysis (PCA) to an expanded longitudinal dataset (n = 2’049 worms, ∼140’000 observations, N_exp_ = 33). Principal component 1 (PC1) explained 24.9% of total phenotypic variance (Fig. S4a). Data points progressively shifted along the PC1 axis with increasing relative age (Fig. 2a), indicating that aging is a major driver of phenotypic variation. Features declining with age showed negative loadings on PC1, whereas those increasing with age showed positive loadings (Fig. S4b). PC1 also captured aging trajectories at the level of individual worms (Fig. 2b). However, its limited predictive accuracy suggested that non-linear feature dynamics contribute substantially to age-related phenotypic variation (Fig. 2c, Fig. S4c-d).

**Fig 2.**
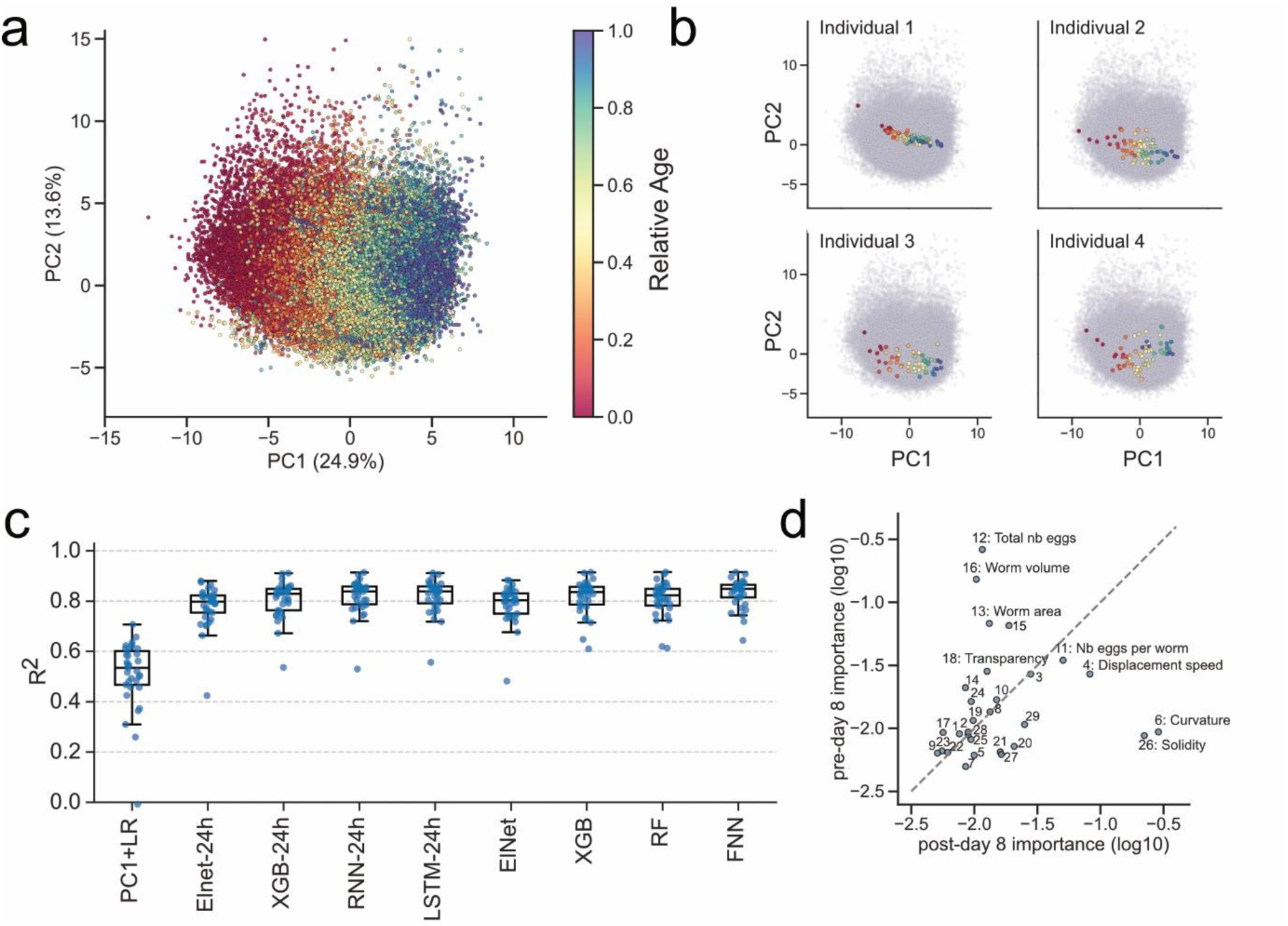
Different machine learning architectures enable accurate quantification of relative age. **a**, Principal component analysis (PCA) of the 141’014 recordings from worms alone in their micro-chamber. Axes show percentage of variance explained by principal components 1 (PC1) and 2 (PC2). Scale bar gives the correspondence between color and relative age. **b**, Examples of individual aging trajectories (colored) for four worms in PCA-space (grey cloud). **c**, Coefficient of determination (R^2^) values in predicting relative age obtained from the different models tested. Models were evaluated with Leave-One-Group-Out Cross-Validation (LOGOCV) with N_exp_ = 33 (see Methods), PC1+LR : PC1 + Linear Regression; Elnet: ElasticNet. XGB: XBoost. RF: Random Forest; RNN: Recurrent Neural Network; LSTM: Long-Short-Term Memory. -24h suffix indicates models that were trained with 4 successive datapoints as input, models without suffix use 1 datapoint as input. Box-plots show the median (central line), interquartile range IQR (boxes), and 1.5 x IQR (whiskers). Individual dots show the performance per experiment. This description of boxplots applies to all following panels. **d**, Comparison of features’ importance (gain) in XGB models trained before day 8 (y-axis) and after day 8 (x-axis). Each point represents one feature. The grey dashed line indicates the line of perfect fit. Features numbering (ordered as in fig 1c): 1: Head ampl.; 2: Midbody ampl.; 3: Tail ampl.; 4: Displacement speed; 5: Body bending freq.; 6: Curvature; 7: Shape 1; 8: Shape 2; 9: Shape 3; 10: Shape 4; 11: Nb eggs per worm; 12: Total nb eggs; 13: Worm area; 14: Worm length; 15: Worm perimeter; 16: Worm volume; 17: Tail width; 18: Transparency; 19: transparency norm.; 20: Intestine area; 21: Intestine width; 22: Intestine area / worm area; 23: Intestine area / tail area; 24: Intestine width / worm width; 25: Intestine width / tail width; 26: Solidity; 27: Solidity SD; 28: Orientation SD; 29: Eccentricity.

We next implemented supervised machine learning (ML) models to predict each worm’s relative age at any given time point. Models were trained using longitudinal phenotypic measurements collected over four consecutive recordings (6h intervals; 24h window), concatenated into a single feature vector, with relative age at the final time point as the target variable. We evaluated four model classes: Elastic Net (ElNet), a regularized linear approach frequently used in aging clocks^29^; eXtreme Gradient Boosting (XGB), a tree-based ensemble method^30^; and two sequence-aware neural network architectures, recurrent neural networks (RNN)^31^ and long short-term memory networks (LSTM). All models trained on 24h windows substantially outperformed the PC1 + linear regression baseline, explaining >80% of the variance in relative age on average (RMSE ∼0.12) (Fig. 2c, Fig. S4d-e, Table S1).

To determine whether temporal context was required for accurate prediction, we trained models using single time-point measurements. Predictive performance remained comparable to that of 24h-windows models (median R² = 0.79-0.85; RMSE = 0.12-0.13) (Fig. 2c, Fig. S4d, Table S1), indicating that multidimensional phenotypic measurements contain sufficient information to estimate relative age across the lifespan. We therefore examined whether the predictive contribution of individual traits varies across life stages. Feature importance analysis revealed stage-specific predictors (see Methods). Early adulthood (0–8 days, encompassing the reproductive period^32^) was primarily characterized by reproductive output, body size metrics (perimeter and volume), and optical transparency, whereas motility contributed minimally (Fig. 2d). In later life (>8 days), movement-related features, including displacement speed, body curvature, and solidity, became dominant predictors.

These results show that diverse supervised ML architectures accurately estimate relative age from phenotypic trajectories, consistently outperforming a PCA-based baseline. Similar performance across linear, tree-based, and neural network models indicates that predictive accuracy primarily reflects the information content of the phenotypic feature set rather than model choice. We selected the RNN trained on 24 h windows for subsequent analyses because it achieved the lowest median RMSE while remaining computationally efficient; we refer to its predictions as *phenotypic age*.

### Divergent phenotypic aging trajectories emerge within an isogenic population

*C. elegans* are self-fertilizing hermaphrodites that generate genetically identical populations. Despite this genetic homogeneity, substantial inter-individual variability in lifespan persists even under tightly controlled environmental conditions^33^. Whether long-lived individuals simply experience delayed aging or instead follow qualitatively distinct aging trajectories remains debated^22,33–36^.To address this question, we stratified the single-worm N2 dataset (from Fig. 1) into three equally sized lifespan groups (short-, intermediate-, and long-lived) and compared phenotypic dynamics between the shortest-lived (4-16.5 days) and longest-lived (19.5-28 days) subgroups. Predicted phenotypic age followed similar trajectories in both groups until approximately day 6 of adulthood (Fig. 3a,b). However, divergence in specific phenotypic traits emerged earlier. As early as day 2 of adulthood, long-lived worms displayed reduced total egg production, smaller absolute intestinal area, and increased body length compared with short-lived individuals (Fig. 3c-e, Fig. S5a). By day 6, additional differences became apparent, including larger intestine area normalized to body size and increased tail movement amplitude (Fig. 3f,g). Phenotypic divergence continued to widen with chronological age (Fig. S5b,c). From day 8 onward, long-lived worms exhibited higher motility, improved late-life posture, and prolonged growth trajectories relative to short-lived animals. These observations indicate that phenotypic differences emerge at distinct chronological ages depending on the physiological system considered.

**Fig. 3.**
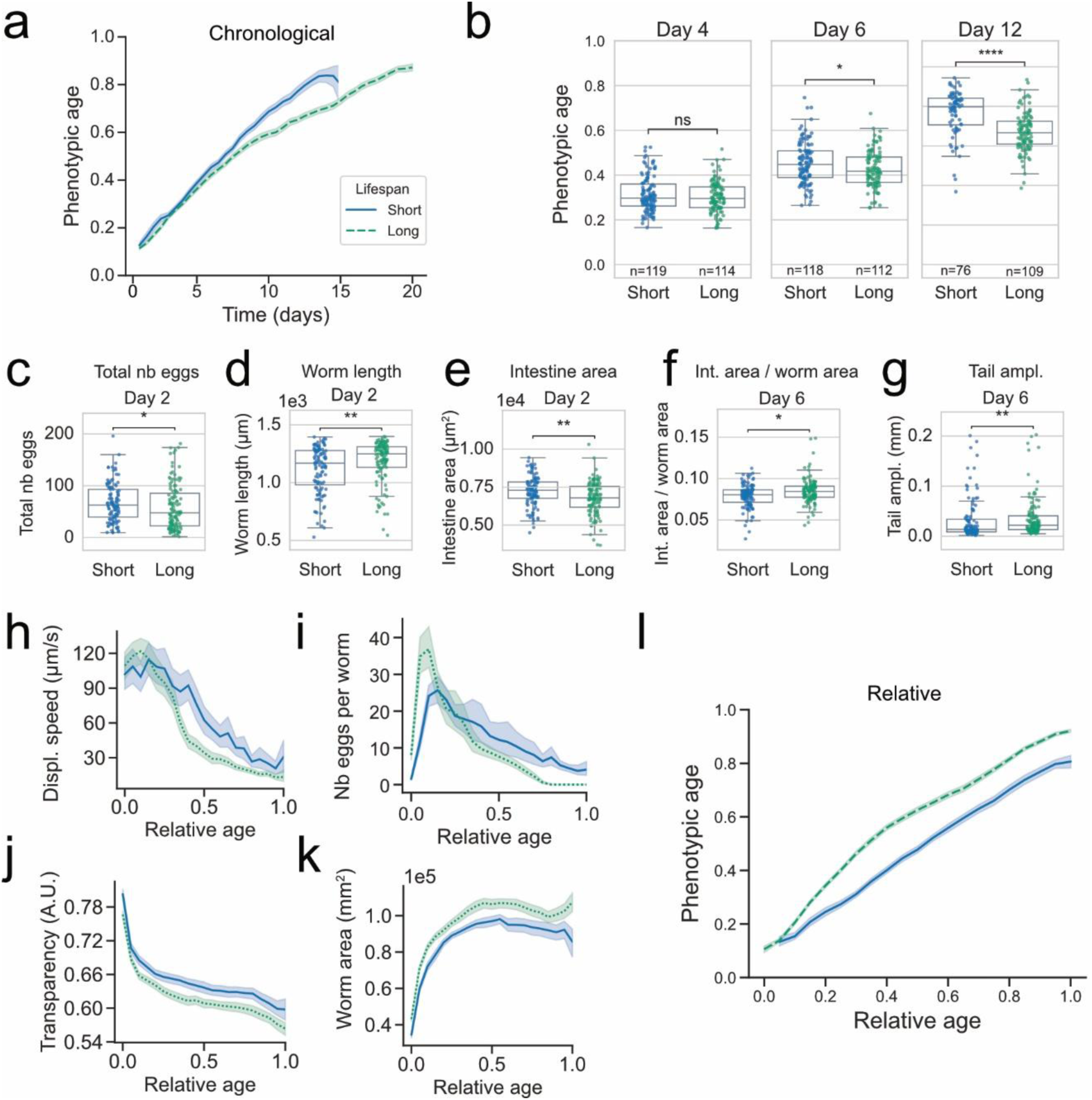
Distinct phenotypic aging trajectories among isogenic wild-type individuals. **a**, Phenotypic age over time in days for the short-lived (blue) and long-lived (green) cohorts (N=122 worms in each). **b**, Phenotypic age distribution within each cohort on different days. Statistical comparisons were performed using two-sided Mann-Whitney U tests. **c-g**, boxplots of features with statistically significant difference before day 7 between short- and long-lived worms. For each feature, comparisons between cohorts were performed every two days using two-sided Mann-Whitney U tests, followed by Benjamini-Hochberg correction for multiple testing (feature-wise). **c**, total number of eggs laid up to day 2, **d**, worm length at day 2, **e**, intestine area (non-normalized) at day 2, **f**, normalized intestine area at day 6, **g**, tail amplitude at day 6. **h-k**, measurements of selected phenotypic features as a function of relative age (fraction of lifespan) in short-lived and long-lived N2 worms. Short-lived worms are shown as solid blue lines and long-lived worms as green dotted lines. **h**, displacement speed, **i**, Number of eggs per worm, **j**, Transparency,**k**, worm area. **l**, Phenotypic age over relative age for the short-lived (blue) and long-lived (green) cohorts. Significance codes: ns, not significant; *, *P* < 0.05; **, *P* < 0.01; ***, *P* < 0.001; ****, *P* < 0.0001. This applies to all following panels.

To determine whether these differences reflect delayed aging or temporal scaling relationships with lifespan, we analyzed phenotypic traits as a function of relative age. Motility-related features (including late-life features), reproduction, transparency, and growth displayed largely synchronized trajectories between lifespan groups when normalized to lifespan (Fig. 3i-l, Fig. S7-S12). Consequently, relative to total lifespan, long-lived worms reproduced over a shorter fraction of life, continued growing over a longer fraction, and remained in low-motility states for longer than short-lived individuals. In contrast, intestine-related traits scaled proportionally with lifespan (Fig. S10). Notably, at equivalent fractions of lifespan, long-lived worms appeared phenotypically older than short-lived worms (Fig. 3l).

These results indicate that phenotypic aging trajectories do not simply scale with lifespan. Rather than exhibiting a uniform delay in aging, long-lived worms experience extended periods of late-life decline relative to their lifespan. These findings support a model in which genetically identical individuals follow distinct trait-specific aging trajectories. Importantly, our phenotypic clock captures these lifespan-associated differences in aging dynamics within an isogenic population.

### Dietary Restriction (DR) modalities converge on a shared phenotypic aging signature

Dietary restriction (DR) is a robust lifespan-extending intervention conserved across taxa^37^. In *C. elegans*, DR can be achieved by reducing bacterial food availability or through genetic perturbations that mimic reduced nutrient intake or activate DR-associated pathways^38,39^. We leveraged three complementary DR paradigms: reduced bacterial density (12.5% of ad libitum; N2 DR), the feeding-defective mutant *eat-2(ad465)*, characterized by reduced pharyngeal pumping^40^, and *slcf-1(tm2258)*, in which loss of the monocarboxylate transporter SLCF-1 activates DR-associated molecular programs^41,42^. These independent modalities allowed us to test whether our phenotypic clock detects intervention-dependent changes in aging trajectories and whether distinct DR paradigms converge on a common phenotypic signature.

Consistent with prior work, DR significantly extended lifespan (Fig. 4a). Wild-type animals exposed to reduced bacterial density from the L4 stage throughout adulthood exhibited a 31% increase in median lifespan and a 46% increase in maximal survival (95th percentile; p < 0.0001) relative to ad libitum (AL) controls (Fig. 4a). Genetic DR models also showed increased longevity, albeit to a lesser extent: median lifespan increased by 21% in *eat-2(ad465)* and 18% in *slcf-1(tm2286)*, while maximal survival increased by 46% and 24%, respectively (Fig. 4a).

**Fig. 4.**
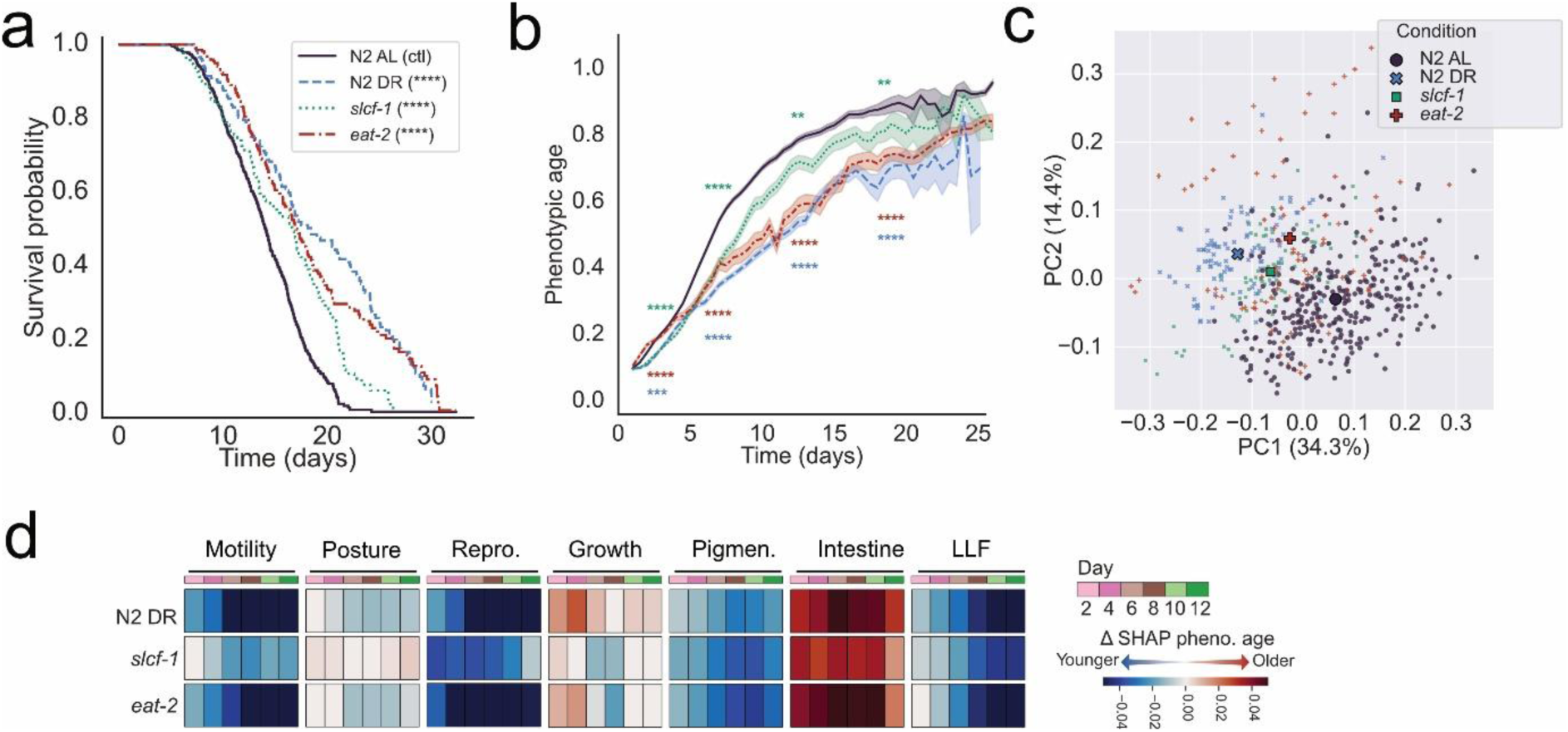
Dietary Restriction (DR) modalities converge on a shared phenotypic aging signature. **a**, Survival curves of N2 AL (*ad libitum*), N2 DR, *eat-2(ad465)*, *slcf-1(tm2286)*. Median survival in days were 14.1, 18.4, 17.1, 16.6, for the AL, DR, *eat-2*, and *slcf-1* conditions respectively. N=487, 143, 182, 109. Statistical significance was assessed using two-sided log-rank (Mantel–Cox) tests compared to AL. Controls of the different experiments were pooled (N_exp_ = 6). **b**, Longitudinal prediction of phenotypic age by the RNN model over 25 days of adulthood. Solid lines represent the mean relative age per condition; shaded areas indicate 95% CI (bootstrap, *n*_boot_= 1’000). Mann–Whitney U tests performed on N2 DR, *eat-*2 and *slcf-1* against AL control at days 2, 6, 12 and 18. *P* values were corrected for multiple comparisons using the Benjamini-Yekutieli procedure. **c**, PCA plot of phenotypic aging trajectories based on SHAP values of the first 12 days of worms from the AL, DR, *slcf-1* and *eat-2* conditions. Centroids are also shown. **d**, Heatmap of differences in SHAP values (ΔSHAP) with controls (fed *ad lib.*) for DR, *eat-2*, and *slcf-*1, derived from the RNN-24h phenotypic aging clock. Non-significant differences (two-sided Mann-Whitney U test) were set to 0.

Our phenotypic clock detected consistent reductions in phenotypic age across DR conditions (Fig. 4b). Wild-type animals subjected to DR displayed younger phenotypic ages than AL controls across most of adulthood, with divergence emerging shortly after intervention onset (Fig. 4b, Fig. S13). Similarly, *eat-2(ad465)* and *slcf-1(tm2258)* mutants exhibited reduced phenotypic age compared with wild-type animals under AL condition. The magnitude and kinetics of phenotypic age reduction varied between modalities, with weaker effects observed in *slcf-1(tm2258)* and delayed divergence in *eat-2(ad465)* relative to AL. These differences paralleled variation in lifespan extension, indicating that the phenotypic clock captures graded intervention effects on organismal aging dynamics.

To determine whether distinct DR paradigms share a common phenotypic signature, we quantified feature-level contributions to phenotypic age using SHAP (SHapley Additive exPlanations) values (see Methods). Principal component analysis of SHAP values revealed clear separation between DR conditions (N2 DR, *eat-2(ad465)*, *slcf-1(tm2258)*) and AL controls (Fig. 4c). Hierarchical clustering confirmed segregation into DR versus non-DR groups (Fig. S14), indicating convergence toward a shared phenotypic space despite mechanistic differences between interventions.

Although DR conditions were globally associated with younger phenotypic age, SHAP analysis revealed trait-specific remodeling rather than uniform rejuvenation (Fig. 4d). Individual traits shifted in both directions relative to AL controls, with some traits adopting younger states and others older states. The amplitude of traits-level changes was smallest in *slcf-1(tm2258)*, consistent with its weaker phenotypic and survival effects. For example, reproductive traits were less strongly altered in *slcf-1(tm2258)* than in *eat-2(ad465)* mutants, in agreement with previous observations ^41^.

Thus, our phenotypic clock robustly captures DR-dependent modulation of aging trajectories across multiple intervention modalities. Distinct DR paradigms converge on a shared phenotypic signature characterized by an overall younger phenotypic age arising from coordinated but trait-specific remodeling of physiological features.

### Distinct phenotypic aging trajectories underlie similar lifespan extension after early- and late-onset DR

The timing of geroprotective interventions can critically influence their impact on aging trajectories, yet whether comparable lifespan extensions arise from similar or distinct organismal states remains unclear. To address this, we implemented four feeding regimens differing in the timing of DR: ad libitum feeding throughout adulthood (AL), lifelong DR at 20% food availability (DR), ad libitum feeding followed by DR initiated on day 6 of adulthood (AL-DR), and early-life DR followed by a return to ad libitum feeding from day 6 onward (DR-AL) (Fig. 5a). Lifelong DR produced the strongest lifespan extension, increasing median lifespan by 49% relative to AL controls (Fig. 5b). In contrast, DR limited to early adulthood (DR-AL) or initiated post-reproduction (AL-DR) resulted in smaller but comparable increases in median lifespan (+18% relative to AL), indicating that distinct intervention schedules can yield similar survival outcomes.

**Fig 5.**
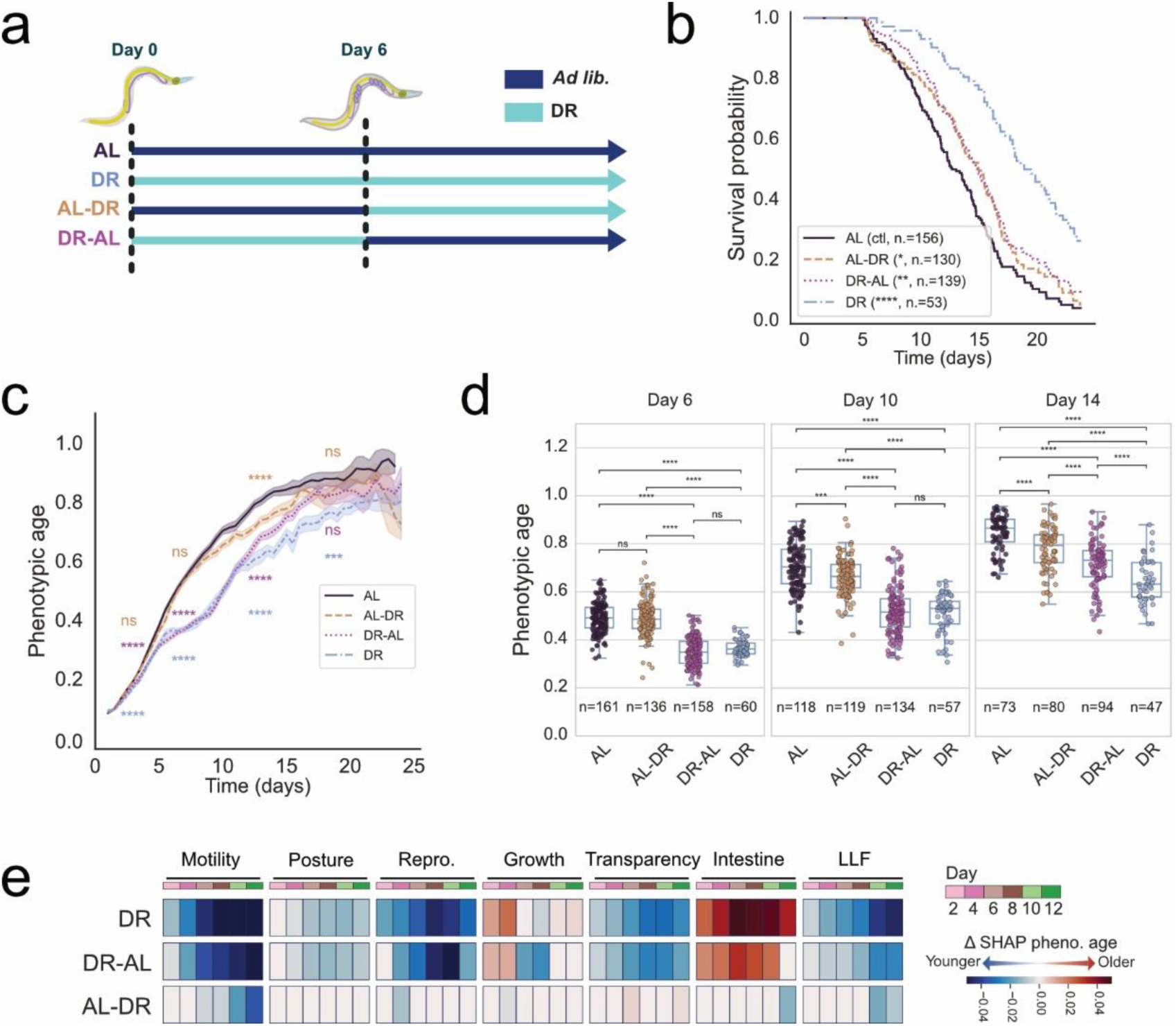
Dietary restriction effects on aging phenotypes vary with the timing of intervention. **a**, Schematic of dietary restriction (DR) protocols. Worms were maintained either on *ad libitum* (AL) or DR conditions throughout life or switched from one diet to another on Day 6. **b**, Kaplan-Meier survival curves of N2 worms under the four dietary conditions. Statistical significance was assessed using two-sided log-rank (Mantel–Cox) tests compared to AL. Median survival times were: 12.7 days (AL), 18.9 days (DR), 15.0 days (AL-DR) and 14.8 days (DR-AL). N = 156, 53, 130, and 139 worms per condition, respectively. **c**, Longitudinal prediction of phenotypic age by the RNN model over 25 days of adulthood. Solid lines represent the mean relative age per condition; shaded areas indicate 95% CI (bootstrap, *n*_boot_= 1’000). Mann–Whitney U tests comparing DR conditions to the AL control at days 2, 6, 12 and 18. *P* values were corrected for multiple comparisons using the Benjamini-Yekutieli procedure. **d**, Phenotypic age for individual worms at days 6, 10 and 14 of adulthood. Points represent individual animals. Two-sided Mann-Whitney U tests with Benjamini-Hochberg correction for multiple comparisons. **e**, Heatmap of ΔSHAP for DR, AL-DR and DR-AL conditions compared to control (AL), derived from the RNN-24h phenotypic aging clock. Non-significant differences (two-sided Mann–Whitney U test) were set to 0.

Analysis of phenotypic age trajectories revealed pronounced differences in organismal responses to DR timing (Fig. 5c,d). Worms exposed to early-life DR (DR-AL) closely followed the phenotypic trajectory of lifelong DR animals until approximately day 12 of adulthood and remained phenotypically younger than AL controls until day 17. In contrast, worms switched from AL to DR at day 6 (AL-DR) exhibited a delayed and transient reduction in phenotypic age, appearing younger than AL controls between days 8 and 16. However, neither phenotypic age trajectories nor SHAP feature profiles converged toward those observed under lifelong DR (Fig. 5c-e), indicating that late-onset DR only partially recapitulates the phenotypic effects of continuous DR.

Together, these findings show that DR reshapes aging trajectories in a time-dependent manner. Notably, early- and late-onset DR produced similar extensions in lifespan despite inducing distinct phenotypic aging dynamics. These results demonstrate that comparable survival outcomes can arise from divergent organismal states, highlighting the importance of longitudinal phenotypic measurements to resolve how intervention timing modulates biological aging trajectories.

### Longitudinal phenotyping reveals time-dependent genetic interactions shaping aging trajectories

We next asked whether longitudinal phenotypic age measurements could resolve epistatic interactions in aging pathways that are not apparent from survival outcomes alone. To address this question, we analyzed *slcf-1(tm2258)* and *daf-18(e1375)* mutants and their corresponding double mutant. In *C. elegans*, *daf-18*, the ortholog of the tumor suppressor PTEN, is required for normal lifespan and contributes to hormetic stress responses induced by *slcf-1* downregulation^41,42^. However, the extent to which *daf-18* contributes to *slcf-1*-mediated longevity remains unclear.

Lifespan measurements obtained in our microfluidic system recapitulated previous observations (Fig. 6a). Loss of *daf-18* shortened lifespan relative to wild type, whereas mutation of *slcf-1* increased median lifespan by ∼20% in both wild-type and *daf-18* mutant backgrounds. These results suggest that lifespan extension conferred by *slcf-1* is largely independent of DAF-18 activity when evaluated at the level of survival.

**Fig. 6.**
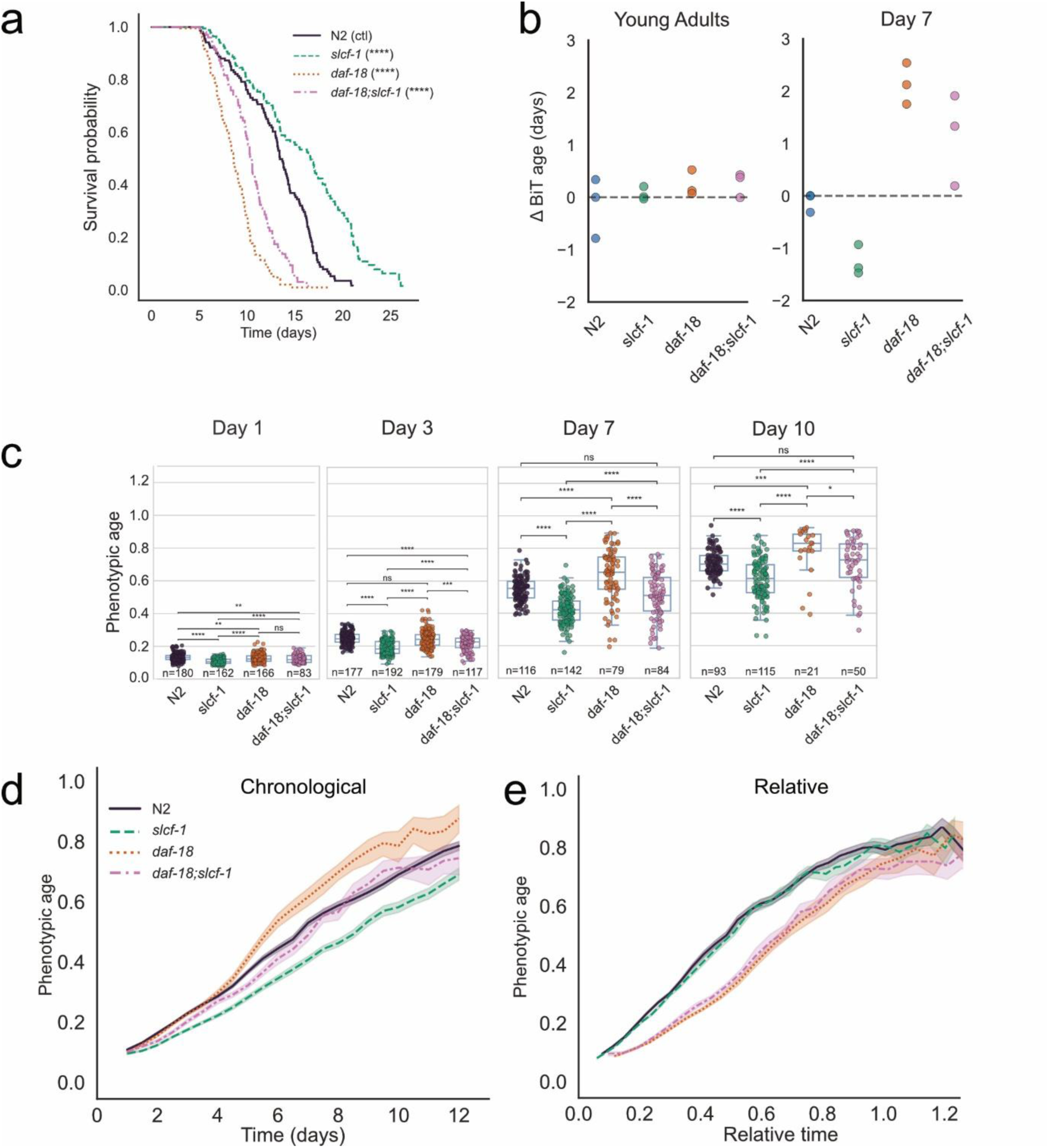
Longitudinal tracking of *daf-18;slcf-1* double mutants suggest an age-dependent effect of *daf-18* on phenotypical aging. **a**, Kaplan–Meier survival curves for N2, *slcf-1(tm2286)*, *daf-18(e1375)*, and *daf-18(e1375)*;*slcf-1(tm2286)* double mutants. Median survival times (in days): N2, 13.4d (N=125); *slcf-1(tm2286)*, 16.3d (N=109); *daf-18(e1375)*, 8.6d (N=104), *daf-18(e1375)*;*slcf-1(tm2286)*, 10.3d (N=82). Statistical significance was assessed using two-sided log-rank (Mantel–Cox) tests compared to N2. **b**, Predicted BiT ΔAge in days, using transcriptomic data from Young Adults and 7-day-old adult worms. **c**, Phenotypic age for individual worms at days 3, 7 and 10. Two-sided Mann-Whitney U tests with Benjamini-Hochberg correction for multiple comparisons. **d**, Predicted phenotypic age by the RNN model over time in days by the RNN-24h. Lines show the condition mean and shaded area represent the 95% CI of the mean (by bootstrap, with n_boot_=1000). **e**, Predicted phenotypic age by the RNN model over relative time by the RNN-24h. Lines show the condition mean and shaded area represent the 95% CI of the mean (by bootstrap, with n_boot_=1000).

To determine whether these mutations differentially influence biological age, we compared two complementary measurements: transcriptomic age and phenotypic age. Transcriptomic age was estimated using the BiT age clock^19^. In young adults, all genotypes exhibited similar transcriptomic ages (Fig. 6b). By day 7, however, *daf-18* mutants were predicted to be biologically older than wild-type animals, whereas *slcf-1* mutants appeared younger. These trends were recapitulated by our phenotypic clock, which detected significant genotype-dependent differences in biological age (Fig. 6c). In double mutants, the two clocks showed partially divergent predictions: transcriptomic age more closely resembled the *daf-18* profile, whereas phenotypic age was intermediate between the two single mutants. These single time-point measurements suggest partial suppression of the *slcf-1* phenotype by *daf-18*, but do not reveal when divergence in aging trajectories emerges.

Longitudinal phenotypic trajectories provided additional temporal resolution (Fig. 6d). Consistent with earlier results, *slcf-1* mutants exhibited slower phenotypic aging than wild-type animals from early adulthood onward. In contrast, *daf-18* mutants initially followed a trajectory similar to wild type until approximately day 5 of adulthood, after which phenotypic aging accelerated. In *daf-18; slcf-1* double mutants, divergence from the *slcf-1* trajectory emerged as early as day 3, earlier than in the wild-type background, and was followed by accelerated phenotypic aging relative to *slcf-1*single mutants. These results indicate an early genetic interaction between *daf-18* and *slcf-1* that alters the temporal dynamics of phenotypic aging. Notably, when trajectories were expressed as a function of relative age, the double mutant aligned with *daf-18* single mutants (Fig. 6e), indicating that *daf-18* phenotype predominates in shaping the aging trajectory of *slcf-1* mutants across the lifespan.

Hence, these analyses show that phenotypic age trajectories broadly correlate with transcriptomic age while providing substantially higher temporal resolution. Although *slcf-1* extends lifespan in a *daf-18* mutant background, DAF-18 is required for the characteristic temporal pattern of *slcf-1*-mediated slowing of aging. More broadly, these results illustrate how longitudinal phenotypic measurements can reveal time-dependent epistatic interactions that are not detectable using lifespan alone

### Phenotypic clock enables early detection of multidimensional effects of candidate geroprotectors

Lifestyle interventions such as dietary restriction (DR) robustly extend lifespan, yet their long-term implementation in humans remains challenging, motivating the identification of pharmacological alternatives (geroprotectors)^43^. However, candidate geroprotective compounds often produce modest lifespan extensions that approach the intrinsic variability of survival measurements, making their evaluation slow, costly, and frequently inconclusive when relying solely on lifespan. We therefore applied our longitudinal phenotyping framework to pharmacological interventions expected to produce subtler effects than the aging mutants described above, the biguanide metformin and the NAD⁺ precursors nicotinamide riboside (NR) and nicotinamide (NAM).

Metformin has been shown to extend lifespan in *C. elegans* and mice^44,45^ and recent work indicates that it can slow transcriptomic aging clocks in non-human primates^3^. Age-associated depletion of nicotinamide adenine dinucleotide (NAD⁺) has been linked to multiple age-related pathologies, including sarcopenia, prompting interest in interventions that restore NAD⁺ homeostasis^46,47^. We first evaluated metformin hydrochloride at 25 mM and 50 mM, concentrations previously reported to increase *C. elegans* lifespan ^44^. Under our experimental conditions, both doses produced modest increases in median lifespan (+13.5% and +8.6%, respectively; Fig. 7a). In contrast, longitudinal phenotypic analysis revealed consistent reductions in phenotypic age under both metformin treatments (Fig. 7b,c). Metformin-treated animals could be distinguished from untreated controls as early as day 4 of adulthood, well before differences in survival outcomes became apparent. The higher concentration produced a more sustained shift toward younger phenotypic age across adulthood. SHAP decomposition indicated that this reduction was primarily driven by improvements in motility, transparency, and reproductive output, whereas intestine-associated traits shifted toward relatively older states, yielding a multidimensional profile partially reminiscent of DR-associated phenotypic remodeling (Fig. 7d and Fig 4d).

**Fig. 7.**
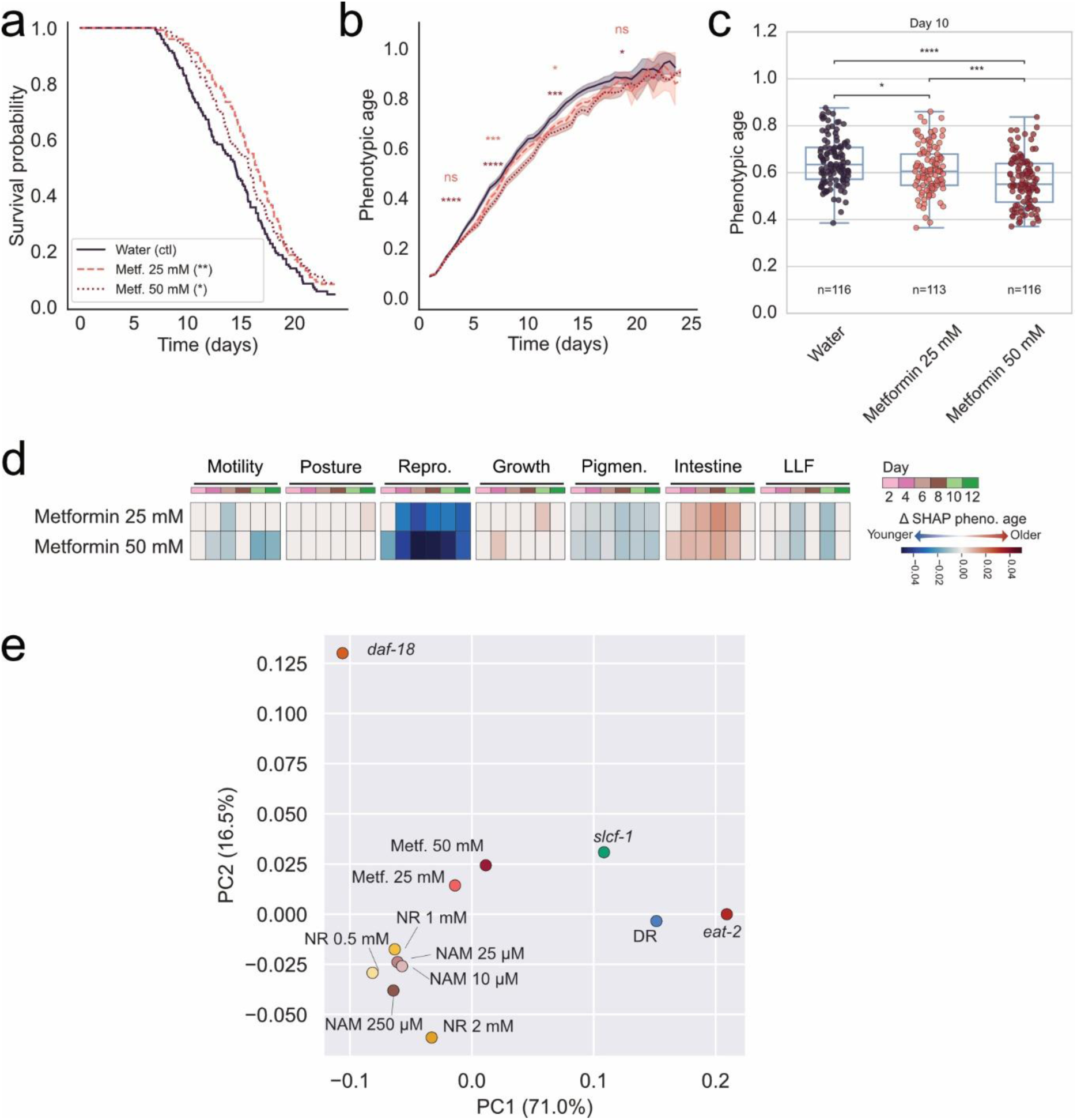
Identification of geroprotectors through a phenotypic clock. **a**, Kaplan-Meier survival curves of N2 worms treated with Metformin at 25 and 50 mM concentrations. Two-sided log-rank (Mantel–Cox) tests. Median survival times were: 14.5 days (untreated), 16.4 days (Met 25 mM), 15.7 days (Met 50 mM). N = 139, 114, 119 worms per condition, respectively. **b**, Longitudinal prediction of phenotypic (relative) age by the RNN model over 25 days of adulthood. Solid lines represent the mean relative age per condition; shaded areas indicate 95% CI (bootstrap, *n*_boot_= 1’000). Two-sided Mann–Whitney U tests against control at days 2, 6, 12 and 18. *P* values were corrected for multiple comparisons using the Benjamini-Yekutieli procedure. **c**, Phenotypic age of metformin-treated worms compared to control N2 (water) for individual worms at day 10. Statistical significance was assessed using two-sided Mann-Whitney U tests with Benjamini-Hochberg correction for multiple comparisons. **d**, Heatmap of differences in SHAP traits values with controls of Metformin at 25 and 50 mM concentrations derived from the RNN-24h phenotypic aging clock. Non-significant differences (two-sided Mann-Whitney U test) were set to 0 in the heatmap. **e**, PCA plot using differences in SHAP traits values compared to respective control conditions.

We next examined NAD⁺-boosting compounds by testing NAM and NR across three concentrations each. None of the tested conditions significantly reduced overall phenotypic age. Nevertheless, higher doses induced selective functional improvements, most prominently increased motility and transparency at 2 mM NR (Fig. S15), demonstrating that longitudinal phenotyping can detect domain-specific physiological effects even in the absence of measurable lifespan extension.

To directly compare pharmacological interventions with established longevity paradigms, we projected SHAP-derived trait differences induced by drug treatments together with those associated with DR conditions (DR, *eat-2*, *slcf-1*) and *daf-18* mutants into a shared phenotypic space using PCA (Fig. 7e). NAD⁺-boosting conditions clustered together, with the highest concentrations (NAM 250 µM and NR 2 mM) producing the strongest phenotypic shifts. Metformin-treated animals localized closer to DR-associated phenotypic states, with the highest tested concentration producing the largest displacement toward the DR region of phenotypic space.

Together, these results demonstrate that longitudinal phenotypic age measurements provide a sensitive framework to evaluate candidate geroprotective compounds beyond survival outcomes alone. By detecting intervention-associated shifts in phenotypic trajectories as early as day 4 of adulthood, this approach enables rapid identification of multidimensional physiological effects even when lifespan changes are modest. Longitudinal phenotyping therefore provides a scalable strategy to prioritize candidate geroprotectors and characterize their functional signatures across the lifespan

## Discussion

Understanding how interventions influence aging requires quantitative frameworks that capture organismal state across time rather than relying solely on survival outcomes. Here we developed a longitudinal phenotyping platform that integrates high-frequency behavioral, morphological, and physiological measurements to quantify multidimensional aging trajectories in *C. elegans*. Across genetic perturbations, dietary interventions, and candidate geroprotective compounds, our results indicate that aging cannot be fully described by a single rate parameter but instead reflects dynamic remodeling of organismal state throughout the lifespan. Distinct lifespan-extending interventions converged on shared phenotypic signatures associated with a younger phenotypic age, yet similar longevity outcomes also arose from divergent aging trajectories depending on intervention timing or genetic background. By resolving these temporal dynamics, longitudinal phenotypic age provides a quantitative framework to dissect how environmental and genetic factors reshape biological aging trajectories and reveals multiple physiological paths to lifespan extension.

Microfluidic technologies have transformed *C. elegans* research by enabling precise environmental control and automated longitudinal measurements under standardized conditions ^48–51^. Using the SydLab™ One platform, we monitored multiple aging biomarkers in parallel at high temporal resolution, generating a high-content atlas of organism-level aging trajectories across the lifespan. Age-associated changes in motility ^33,52,53^, behavior ^22,34,54^, reproduction ^32,55^, body size ^8,31^, transparency ^27,56^, intestinal morphology ^57–59^, and image-derived morphological descriptors ^22^ recapitulated and extended previous descriptions of physiological decline in *C. elegans* ^33,53,60^. By capturing these features simultaneously within individuals, this framework enables direct comparison of aging dynamics across physiological systems.

Leveraging this dataset, we developed MOSAIC, a non-invasive video-based phenotypic clock capable of estimating relative age from short (2s) recordings obtained at any stage of adulthood. Multiple machine learning architectures achieved similar predictive performance, indicating that accuracy primarily derives from the biological depth of the phenotypic features rather than the modeling approach itself. The contribution of individual traits to phenotypic age prediction varied across the lifespan, underscoring the context-dependent nature of aging biomarkers. Compared with previous phenotypic predictors of biological age ^22,25,35^, the improved performance of our approach likely stems from the combination of high sampling frequency, large cohort size, and multidimensional feature integration. Phenotypic age estimates correlated with transcriptomic age predicted by the BiT age clock, supporting the biological relevance of this non-invasive measure. Because it relies exclusively on bright-field imaging, this strategy enables repeated measurements in the same individuals and provides a scalable framework to quantify how interventions dynamically modulate biological aging trajectories.

Our results further indicate that biological age is inherently multidimensional and trait-specific. While temporal scaling of survival curves has been interpreted as evidence that longevity interventions act on shared determinants of aging, lifespan extension can also occur alongside selective preservation or deterioration of specific physiological functions ^7,8,25,34,36,61–64^. Consistent with this view, long-lived wild-type animals appeared phenotypically younger than short-lived individuals of the same chronological age. Yet, these long-lived individuals also appeared older when phenotypic age was expressed relative to lifespan, consistent with an extended period of late-life frailty described previously ^35^. Several traits exhibited similar timing of decline across lifespan groups, whereas others, including late-life motility and behavioral features, were selectively preserved in long-lived individuals. Intestinal traits showed the strongest temporal scaling with lifespan, in line with evidence that intestinal deterioration is a major proximate cause of death in *C. elegans*^58,65–67^.Collectively, these findings indicate that biological age emerges from partially independent physiological trajectories rather than a single coordinated decline.

Analysis of canonical longevity interventions further revealed substantial heterogeneity in the structure of aging trajectories. Some perturbations, including DR, *slcf-1* mutation and metformin treatment, primarily reduced the rate of phenotypic aging without markedly altering trajectory shape. By contrast, *daf-18* mutation produced a qualitatively distinct trajectory characterized by accelerated late-life decline. Moreover, comparable lifespan extension could arise from distinct phenotypic histories, as illustrated by early- and late-onset DR paradigms. These observations indicate that interventions influence not only how fast organisms age but also how aging unfolds across physiological systems. Intervention timing emerged as a key determinant of these dynamics, consistent with previous studies showing that modulation of IIS signaling in adulthood can improve late-life traits while minimizing early-life trade-offs ^67,68^.

Identification of geroprotective interventions remains a central objective of geroscience. While *C. elegans* provides a powerful platform for large-scale screening ^50^, lifespan alone incompletely captures intervention effects on organismal function. Indeed, some interventions extend lifespan while impairing physiological performance, whereas others improve functional traits without producing large survival effects ^7,8^. Our phenotypic clock enables evaluation of candidate geroprotectors based on their multidimensional impact on health-related traits, without requiring invasive molecular profiling. Because phenotypic age can be assessed longitudinally and early in adulthood, this approach may substantially accelerate screening workflows by enabling early prioritization of compounds with beneficial effects on organismal function. Furthermore, SHAP-based decomposition of phenotypic age provides insight into how individual physiological systems contribute to intervention effects, facilitating identification of compounds that promote desirable healthspan profiles.

In summary, MOSAIC demonstrates that longevity interventions do more than extend lifespan; they redirect aging along alternative phenotypic trajectories whose structure depends on intervention type, genetic background and timing of exposure. These findings support a model in which biological aging is plastic, multidimensional and shaped by dynamic physiological trade-offs. Longitudinal phenotyping approaches that resolve these trajectories may therefore provide a powerful strategy to identify interventions that optimize both lifespan and healthspan, and to anticipate context-dependent effects of candidate geroprotective therapies.

## Material and Methods

### Buffers preparation

S-medium was prepared by supplementing S-basal (97.32 mM NaCl, 5.74 mM K_2_HPO_4_, 44.08 mM KH_2_PO_4_) with 3.00 mM CaCl_2_, 3.00 mM MgSO4, 10.0 mM potassium citrate, trace metals solution (50 µM EDTA, 24.8 µM Fe^2+^, 10.1 µM Mn^2+^, 10.1 µM Zn^2+^, 1 µM Cu^2+^), 12.93 µM cholesterol, 118 µM carbenicillin, 134 µM ampicillin, 0.06 % (v/v) TWEEN® 20, and 0.195 µM doxycycline. The solution was sterile-filtered through a 0.22 µm membrane before use.

### *C. elegans* strains and culture

The following *C. elegans* strains were used: N2 Bristol, *eat-2(ad465)* II ^40^, *daf-18(e1375)* IV ^69^. *slcf-1(tm2258)* and *slcf-1(tm2258);daf-18(e1375)* come from ^41^.

Strains were maintained on NGM plates seeded with *E. coli* OP50 according to standard protocols (Brenner 1974) at 20°C. All experiments were executed on the SydLab^TM^ One devices set at 20°C starting with synchronized populations of L4 worms.

### Nematodes synchronization and loading into the device

Worm populations were synchronized by egg laying in liquid culture followed by recovery of arrested L1 larvae using a filter-based method and transferred to OP50-seeded NGM plates until the L4 stage.

Synchronized L4 worms were washed in S-medium, diluted to the appropriate concentration, and loaded into the reservoirs of the SydLabTM One platform. At the onset of experiments, synchronized L4 worms were aspirated from the reservoirs and distributed automatically inside the microfluidics chambers, where passive hydrodynamics confined them for the duration of the experiment. The device maintained a constant temperature (20°C) and delivered standardized bacterial suspensions at defined hourly intervals for the duration of the experiments. Feeding, compound exposure, and imaging were then performed automatically by the platform.

### Bacteria preparation

Freeze-dried *E. coli* OP50 were resuspended in S-medium and diluted to a final concentration of approximately 62.5 mg/ml based on the mean OD_600_ to standardize the *ad libitum* feeding condition. For dietary restriction (DR), this preparation was further diluted in S-medium. A fresh bacteria preparation was provided every 2 to 3 days.

### Chemicals preparation

Nicotinamide riboside (NR; Cayman Chemical #36941) was dissolved in deionized water at 100 mM. Metformin hydrochloride (Sigma-Aldrich #317240) was dissolved in deionized water to generate a 0.5 M stock solution. Nicotinamide (NAM; Sigma-Aldrich #N0636) was dissolved in deionized water at 100 mM.

Working concentrations were obtained by direct dilution of stock solutions into the bacterial suspension immediately prior to loading into the microfluidic platform.

### Transcriptomic analyses

We extracted total RNA from three independent replicates of each genotype (wild-type N2, *daf-18(e1375)*, *slcf-1(tm2258)*, and *daf-18(e1375); slcf-1(tm2258)*) at the young adult stage (YA) and on day seven of adulthood (A7). We amplified 100 ng of total RNA with the GeneChip 3 IVT Express kit and hybridized 15 µg of cRNA to Affymetrix *C. elegans* Genome Arrays. We normalized CEL files with the MAS5 algorithm in the Affymetrix Expression Console and filtered expression values in Partek Genomic Suite v6.6 with a Student t test (P < 0.05) and a two-fold change threshold.

Next, we mapped probe sets to WormBase gene identifiers with the *celegans.db* package in 𝑅, averaged duplicate probes, and applied quantile normalization with *limma*. We analyzed this matrix with the BiT-age transcriptomic clock, assigning zero to genes that were missing from the array in every sample. Finally, the predicted BiT ages were rescaled relative to the median of N2 controls at YA stage and seven days for A7.

### Image acquisition and object detection

Brightfield (BF) videos of 2 seconds with 5 frames per second (8-bit, PNG) were acquired for each chamber every 6 hours using a custom-made OEM-microscope equipped with CMOS 4MP monochromatic camera (2048 x 2048px sensor) (Basler), a fluor 10x/0.5NA objective (Zeiss) as well as a custom-made LED panel. Magnification 6x final. Imaging was performed automatically 30 seconds after liquid injection.

Deep learning (DL)-based instance segmentation was performed using a multi-class Mask R-CNN on images to detect worms throughout their entire life as well as their progeny (eggs and larvae). The tail intestine was detected using a fine-tuned version of the worm detection Mask R-CNN.

### Annotation of death & computation of relative age

For the aging clock training set, time of death was manually labelled for 2839 (75.3%) worms, whereas the remaining 923 (24.7%) deaths were annotated using a DL algorithm capable of detecting death events. Time of death in the validation experiments were annotated with the death-detection DL algorithm. In manually annotated experiments, worms that died from internal hatching or vulval rupture were censored and excluded from relative-age analyses. In experiments annotated automatically, deaths detected before day 5 of adulthood were treated as censored.

The DL death detection algorithm works in two step. First, videos are analyzed with a model of the CoAtNet-1 architecture, trained on 41’337 BF videos, which classifies each worm in every frame as “alive” or “dead”. In the second step, a rule-based algorithm aggregates the classifications by chamber and time, identifying worm deaths by monitoring how the total number of live worms decreases over time. A consistent drop in the live count is interpreted as a death.

Relative age was computed per worm by dividing the time of acquisition of a recording by the time of death.

### Motility features

The mathematics behind motility analyses was adapted from ^70^. In this framework, the worm’s body is represented as a one-branch, 100-point skeleton, where each point corresponds to an evenly spaced position along the worm’s body from head to tail.

#### Curvature and Body bending frequency

Worm curvature is calculated by computing the curvature in each point of the worm skeleton and averaging it. The body bending frequency of a worm is obtained by averaging the dominant frequency after fast Fourier transform of the worm’s curvature for the skeleton nodes 20 to 80, in increments of 5.

#### Amplitude of the head, midbody and tail

Head is detected as the most curved extremity of the worm mask, whereas tail is detected as the most pointy extremity of the worm mask. First, the absolute amplitude (*A_abs_*) at each skeleton point is calculated as:

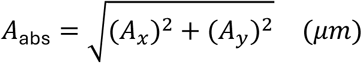

Where:

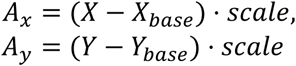

represent the deviation from the baseline in the *x-* and *y-*directions, respectively. *X* and *Y* are the *x-* and *y-*coordinates of the skeleton points over time, and *X_base_* and *Y_base_* are the smoothed baseline coordinates obtained from a polynomial fit of degree 5 over the 10 frames. *Scale* transform pixels to micrometers and is a factor of 1.81.

The head amplitude (*A_head_*) is derived from the first two points of the 100-point skeleton. It is computed as:

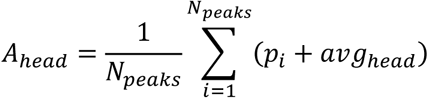

Where:

- *p_i_* are the peak values of ampl abs detected in the head region,
- avg_head_ is the mean amplitude of *A_abs_* in the head region over the 10 frames, and
- *N_peaks_* is the number of peaks detected in the amplitude trace of the head region. Peak detection is performed using a peak-finding algorithm (*find_peaks* from *Scipy signal*), applied to the amplitude trace over the 2-second interval. The final head amplitude represents the mean peak amplitude adjusted by the average baseline amplitude in the head region.

The tail amplitude and mid-body amplitude are calculated in the exact same manner but using the last 2 points of the skeleton and the 3 central points of the skeleton, respectively.

#### Displacement speed

Worm displacement speed (*V_c_*) represents the distance traveled over all frames of the time-lapse video, divided by video duration and is calculated based on the xy position of the centroid of the skeleton as follows:

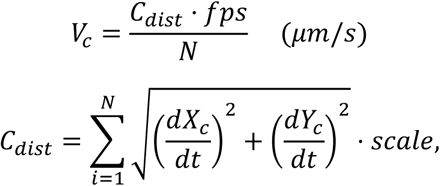

Where:

- *N* is the number of frames in the video,
- 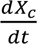 and 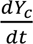 are the time derivatives of the centroid coordinates in the *x-* and *y-*directions, respectively. The centroid coordinates are obtained from a polynomial fit of degree 5 of the skeleton over the 10 frames,
- scale converts distances from pixels to micrometers
- fps is the frame rate of the video (frame per second)

### Posture (shape) analysis

Shape analysis is loosely based on ^54^. In brief, we grouped postures into four broad classes based on (i) how much the body bends overall and (ii) how different the head and tail orientations are. For each frame, from the tangent-angle profile 𝜃(𝑠) we computed:

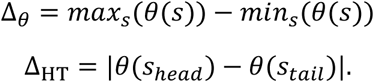

We then assigned a class using fixed radian thresholds:

- **Shape 1** (Nearly straight): Δ_𝜃_ < 𝜋/2.
- **Shape 2**, Single bend (C-like): Δ_𝜃_ ≥ 𝜋/2 and Δ_HT_ < 3𝜋/4.
- **Shape 3** Alternating bend (S-like): Δ_𝜃_ ≥ 𝜋/2 and Δ_HT_ ≥ 3𝜋/4.
- **Shape 4** coiled: Δ_𝜃_ > 3𝜋/2.

If no valid angle trace was available for a frame, it was left unclassified.

### Eggs

Eggs are detected as single eggs or clumps of eggs by the Mask R-CNN model. Therefore, egg counts are performed by dividing the area of the mask with the area of a single egg measured for the N2 strain (1500 pixels). *Number of eggs per worm* amounts to the number of eggs detected in a video at a time *t*, divided by the number of worms in the chamber. *Total number of eggs* is the cumulative sum of *Number of eggs per worm*.

### Worm length, area, perimeter and volume

Similar to motility analysis, skeleton-based analysis is used to compute worm length, by summing the number of pixels forming the skeleton. Worm volume is computed as follow:

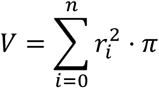

where *n* is the total number of pixels in the skeleton and *r* is the radius given by the distance between each point of the skeleton and the closest edge of the mask. *Worm area* provides the approximate 2D size of a worm by extracting the number of pixels from the detection mask. *Worm perimeter* is calculated using the *scikit-image (v0.22)* function *extract_regionprops*. The *Worm tail diameter* is calculated as follow:

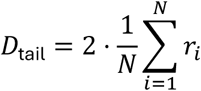

where 𝑟_𝑖_ is the distance from the 𝑖-th skeleton point (within the bottom third of the worm) to the closest edge of the mask, and 𝑁 is the total number of skeleton points in that region.

### Pigmentation

Pigmentation (*Transparency*) is calculated as the average intensity of pixel (values ranging from 0 to 255) inside the worm detection mask divided by the average intensity of pixel outside the mask (*i.e.,* the intensity of the micro-chamber). *Transparency normalized* represent transparency normalized to the mean of the 3 first recordings of *Transparency* for the chamber.

### Intestine

*Intestine area* is the area of the detected tail intestine of worms on the first frame of the video. *Intestine diameter* is the average diameter of the detected tail intestine of worms on the first frame of the video. *Intestine area / worm area, intestine area / tail, intestine diameter / worm diameter* and *intestine area / tail diameter area* are the ratio between the size of the worm and its intestine.

### LLF

The features of the group “Low-Level Features” are derived from the *scikit-image*^71^ function *extract_regionprops*. *Eccentricity* is average eccentricity over the frames of a video. Eccentricity represents how round is an ellipse, with 0 being a circle. *Orientation SD* is the standard deviation of the orientation of the worm over the frames of a video. *Solidity* and *Solidity SD* are the average standard deviation of the *solidity* of the worm over the frames of a video. Solidity is the measure of object compactness and is calculated by dividing the worm mask area with the area of its convex hull.

### Prediction of relative age from individual features

To assess the predictive value of individual phenotypic features, we fitted separate spline-based regression models for each feature. For each model, the target variable was relative age, and the predictor was a single log-scaled phenotypic feature measured at one timepoint. Nonlinear relationships were modeled using a cubic spline basis (*Scikit-learn (v1.3)*^72^ *SplineTransformer*, degree = 3, n_knots = 5, no bias term), followed by z-standardization and ridge regression with internal selection of the regularization parameter over a logarithmically spaced grid of alpha values (*Scikit-learn RidgeCV*; 20 values from 10^-3^ to 10^3^). Model performance was evaluated using 5-fold grouped cross-validation (*Scikit-learn GroupKFold*), with experiments used as the grouping variable to ensure that observations from the same experiment were not split across training and test sets. For each feature, predictive performance was quantified in each test fold using the coefficient of determination (R²), root mean squared error (RMSE) and mean absolute error (MAE), and reported as the mean ± standard deviation across folds.

### Pre-processing, imputation and scaling of longitudinal features for ML algorithms

Occasional gaps in the longitudinal feature tables arose from segmentation failures such as worms momentarily leaving the focal plane, brief losses of microscope focus or misdetection by the mask R-CNN. These missing values were reconstructed in two steps. First, for motility-related variables (head, mid-body and tail bending amplitudes, displacement speed, body-bend frequency and curvature) were interpolated with Akima cubic splines (*Scipy (v1.11)* ^73^ implementation). All remaining variables were interpolated with first-order linear interpolation.

The complete feature matrix was subsequently transformed with a logarithmic function (𝑙𝑜𝑔(𝑥 + 1𝑒^−5^)) as a variance-stabilizing transformation. To attenuate short-term noise while retaining long-term drifts in features, we applied exponentially weighted mean smoothing (smoothing coefficient 𝛼 = 1/3). Last, data were *z*-standardized. All parameters of the *z*-standardization (*StandardScaler* of *Scikit-learn*) were fitted exclusively on the training folds and then re-used to transform test data.

### Construction of ML inputs

For models that operate on one timepoint data (ElasticNet, XGBoost, Random Forest and the feed-forward neural network), each acquisition was transformed to a vector containing the 29 raw features and 3 interaction terms (dynamic power, number of eggs x worm area and volume x darkening), yielding an input vector of dimension (1, 32). Recurrent models (RNN and long-short-term memory (LSTM) networks) received sequences of successive acquisitions; a temporal context window of four time-points (24 h) was used, giving input tensors of shape (4, 32). When sequence-based inputs were incompatible with the chosen model (*e.g.*, ElasticNet, XGBoost), such tensors were reshaped to a single vector of length (1, 128).

### Datasets used for ML development

The *single-worm* dataset comprised 2’053 animals housed individually in micro-chambers, providing 141’014 observations after quality control and exclusion of animals that died from accidental causes (internal hatching or vulval rupture) or were still alive at experiment termination. The *multi-worm* dataset was derived from 800 chambers that contained two or three worms that died within an interval of 72h (1’713 animals in total). For these multi-worm chambers, we generated “synthetic” single-worm records by bootstrapping within each chamber and created 3 synthetic worms per chamber, resulting in the addition of 2453 bootstrapped worms. We did this by sampling half or a third, for the cases of two and three worms, respectively, of the chamber’s recordings with replacement, assigning a synthetic identifier, and defining the time of death by drawing uniformly between the minimum and maximum death times observed in that chamber. After the same interpolation and transformation steps described above, the multi-worm dataset contributed a further 119’741 observations, for a total of 260’755 observations.

### Data augmentation by mix-up

To enlarge the training distribution, we generated virtual samples with two complementary strategies: λ-mixup and feature-mixup. We used both procedures exclusively on the training folds during LOGOCV.

λ-mixup: Each synthetic observation is a convex combination of two randomly selected recordings *a* and *b*. A mixing coefficient λ is drawn from a symmetric Beta distribution *B*(𝛼, 𝛼) (0 ≤ 𝛼 ≤ 2). The feature vector is formed as

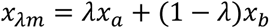

and the corresponding target (relative age) is blended identically,

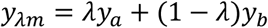

The hyper-parameter 𝛼 controls the expected distance of the synthetic point from its parents; 𝛼 → 0 yields almost exact replicas of real worms, whereas 𝛼 ≈ 2 produces samples centered near the barycenter of the originals. We treated 𝛼 as a tunable parameter and searched the range 0.0-2.0 during hyper-parameter optimization with *Optuna (v3.5)* ^74^, alongside the fraction *p* of additional λ-mixup samples (0-30 % of the training set).

Feature-mixup: This augmentation disrupts the correlation structure at the level of individual variables rather than whole vectors. For every synthetic worm we first sample two donors, *a* and *b*, as above. A Bernoulli mask *m* ∈ {0,1}*^d^* (with *d* = 32 features) selects, independently for each feature, whether the value is taken from donor *a* (*m* = 0) or *b* (*m* = 1). The label is then assigned by weighting the two ages according to the proportion of features originating from each worm, as follows:

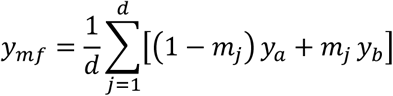

Because feature-mixup does not require a continuous λ, the only free parameter is the proportion *p* of augmented samples; this was optimized over the interval 0-30 %.

### Hyperparameters optimization and training of machine learning models

To quantify biological age we benchmarked five algorithmic families that span linear, tree-based and neural architectures: (i) an *elastic-net* linear model implemented as a stochastic-gradient descent regressor (*Scikit-learn SGDRegressor, penalty = elasticnet*), (ii) *random forests* (*Scikit-learn RandomForestRegressor*), (iii) gradient-boosted decision trees with the *XGBoost (v1.7)* library ^30^, (v) feed-forward neural networks (*FNN*), and (v) recurrent networks (*RNN* and *LSTM*) implemented in *PyTorch (v2.5)* ^75^. Because performance depends critically on the choice of meta-parameters that are not learned during training, we carried out a systematic optimization with *Optuna*, which uses Bayesian optimization (Tree-Parzen estimator). We minimized the root mean squared error (RMSE) of a model.

For each algorithm, we defined a problem-specific search space. These optimizable hyper-parameters consisted of model-specific hyperparameters, two data-augmentation parameters (proportion of feature-mixup and λ-mixup samples; and the β-distribution shape parameter 𝛼; see “Synthetic augmentation by mix-up”), and the sampling ratio applied to the training dataset.

To keep the search computationally reasonable, we evaluated each *Optuna* trial with a five-fold non-stratified cross-validation (CV) that distributed the experiments in folds of approximately equal size – a compromise that reduces the computational burden by a factor of ∼6.5 relative to per-experiment leave-one-group-out CV (LOGOCV). 100 trials were executed per model.

After convergence, the best hyper-parameter configuration was refitted on the full training fraction of each LOGOCV split (33 splits total) and assessed on the held-out experiment; this two-stage procedure yielded an unbiased estimate of generalization while avoiding the prohibitive cost of nested LOGOCV within the *Optuna* optimizer. The predictive performance reported in the “Results” section and Table S1 correspond to models trained with these optimized hyperparameters and evaluated by LOGOCV.

### Age-phase-specific feature importance

To assess whether different features drive predictions early versus late in life, we partitioned the multi-worm dataset by chronological recording day into “early” (days 0–8 of adulthood, inclusive) and “late” (after day 8) subsets. Within each subset, we retrained an XGB model using the same input features, preprocessing, and optimized hyperparameters with *Optuna* of each model as describe above. We extracted gain-based importances for each phase and compared them on a log_10_ scale.

### Prediction with machine learning models

For inferences on unseen experiments, a serialized copy of the best model (trained architecture + hyper-parameters) trained on the complete dataset was loaded and used to predict the relative age of samples. When the experiment to be predicted was part of the original training set, we first retrained an otherwise identical model after removing from the training set every observation from that experiment before predicting, to prevent possibility of information leakage between training and tested data.

Chambers with up to 4 worms were used for inference; for chambers containing more than one worm, we generated *n* bootstrapped worms, equal to the number of worms in the chamber. If a worm died in the interval of time used for the prediction, we removed the data of worms marked as dead by our death detection algorithm, meaning that at any time, there is roughly the same number of predicted worms as the number of alive worms.

Each model returns a continuous estimate 𝑟^(𝑡)of the worm’s *relative age* at chronological time t, where 0≤𝑟^(𝑡)≤1.05. Algorithms with unbounded output (*e.g.*, the ElasticNet regressor) occasionally produced values slightly outside that interval; these estimates were clipped to the nearest bound (0 or 1.05).

### SHAP values

We quantified per-sample feature contribution using the Python Shapley additive explanations (*SHAP (v0.47)*) library ^76^ on the RNN predictions. The RNN and the z-standard scaled training data were passed to a *GradientExplainer*, and this *GradientExplainer* object was used to compute SHAP values of the newly predicted samples.

For the UMAP, a vector containing the aggregated SHAP values from days 2, 4, 6, 8, 10, 12 was created per worm. Worms that died in the interval were excluded from the analysis (because incomplete data). The *ComBat* (*pp.combat*) algorithm from the S*canpy (v1.10)* ^77^ library was used to correct batch effect. UMAP coordinates were computed with the function *pp.neighbors*, followed by calling the *tl.umap* of *Scanpy*. The hierarchical clustering was performed by computing the median SHAP values for each feature, day and condition. These values were transformed to a Z-score across conditions and used as input for hierarchical clustering and visualization with the *clustermap* function from the *Seaborn (v0.13)* ^78^ library.For the SHAP heatmaps, SHAP values were first obtained at the level of the 128 input variables (4 timepoints x 32 features) that enter the network. For ease of interpretation, we subsequently summed the SHAP values to the seven pre-defined functional groups (motility, posture, reproduction, growth, pigmentation, intestine, and low-level features). Values were then binned into 12-h intervals (half-day resolution) according to the chronological time of the recording. Within every bin, we subtracted the median SHAP vector of the appropriate control. Statistical significance was assessed with two-sided unpaired Mann-Whitney U tests were computed compared to control, and non-significant cells were set to 0.

### Survival Analysis

Kaplan-Meier survival analyses were performed using the *lifelines (v0.28)* ^79^ library. Statistical differences between survival distributions were assessed using two-sided log-rank (Mantel-Cox) tests implemented in *lifelines*.

### Longitudinal analysis of phenotypic features

For descriptive trajectories, the 29 phenotypic features were averaged per worm per day and plotted as a function of chronological or relative age.

Inter-feature correlations and correlations between features and relative age were computed after log-transformation of feature values using Spearman’s rank correlation coefficient (*SciPy, scipy.stats.spearmanr*).

For comparisons between groups across multiple time points (e.g., short- and long-lived), two-sided Mann–Whitney U tests were performed at each time point (*SciPy, scipy.stats.mannwhitneyu*). P values were corrected for multiple testing using the Benjamini–Hochberg false discovery rate (FDR) procedure implemented in *stats.multitest.multipletests* function of *statsmodel (v0.14)*.

### GAMM fitting for short- and long-lived cohorts

To distinguish chronologically synchronized features from lifespan-scaled ones we fitted generalized additive mixed models (GAMMs) as a function of both chronological and relative age. These models capture non-linear age-dependent changes while accounting for repeated measurements within individuals. GAMMs were fitted separately for each of the 29 phenotypic features using the *bam* function from the *mgcv R* package with restricted maximum likelihood (REML) estimation. In both analyses, lifespan group was included as a categorical fixed effect (short-lived worms as the reference level), and worm identity was included as a random effect to account for repeated measurements within individuals. To assess differences in trajectory shape over relative age, models were fitted as feature ∼ s(relative_age, by = Lifespan) + Lifespan + s(ID, bs = ’re’), in the syntax of the mgcv package. To assess differences over chronological age, the same formula was used with chronological time in days in place of relative age. For each feature’s model, we extracted the coefficient and associated *P* value for the lifespan-group term from the model summary. P values were corrected for multiple testing using the Benjamini–Hochberg false discovery rate (FDR) procedure implemented in *stats.multitest.multipletests* function of *statsmodel*.

### Features comparisons at specific time points

For analyses restricted to predefined time points (e.g., Fig. S3, Fig. S4), two-sided Mann-Whitney U tests were performed between control and experimental conditions (*SciPy*). When multiple features or time points were tested simultaneously, P values were adjusted using the Benjamini-Hochberg procedure.

### Phenotypic clock age analyses

Differences in predicted phenotypic age trajectories between conditions were assessed using two-sided Mann-Whitney U tests at each time point. To control for multiple comparisons across time points, P values were adjusted using the Benjamini-Yekutieli procedure when dependency between tests was expected.

For comparisons of phenotypic age at specific time points two-sided Mann–Whitney U tests were performed with Benjamini-Hochberg correction for multiple testing. Statistical annotations on plots were generated using the *statannotations* library (relies on *SciPy* for statistical testing).

### Figures creation

Plots were generated using the *matplotlib (v3.8)* ^80^ and *Seaborn* libraries and exported as .svg files. Figures were assembled using Affinity Publisher.

## Authors’ contributions

A.P.V., M.P., F.S. and L.M. designed research; A.P.V., M.P., L.V., M.M., C.N., M.Ch and M.B. performed research; A.P.V., L.S, and F.S. contributed new reagents/analytic tools; A.P.V. and M.P. analyzed data; A.P.V., M.P. and F.S. interpreted data; A.P.V and F.S. wrote the manuscript; A.P.V., M.P. L.V., M.M., M.Co., F.S., and L.M. revised and approved the final manuscript.

## Acknowledgements

The authors thank the ProfileXpert genomics platform (Lyon, France) for sequencing support and technical assistance, Ophélie Muller for assisting with worm maintenance, Youssef Mehdi Attia, Dr. Mirko Francesconi and Pr. Jean-Louis Bessereau for critical discussions. N2 strain was provided by the CGC, which is funded by NIH Office of Research Infrastructure Programs (P40 OD010440). Florence Solari is funded by ANR, grant number ANR-21-CE14-0026-01.

## Supplementary Data

**Fig. S1.**
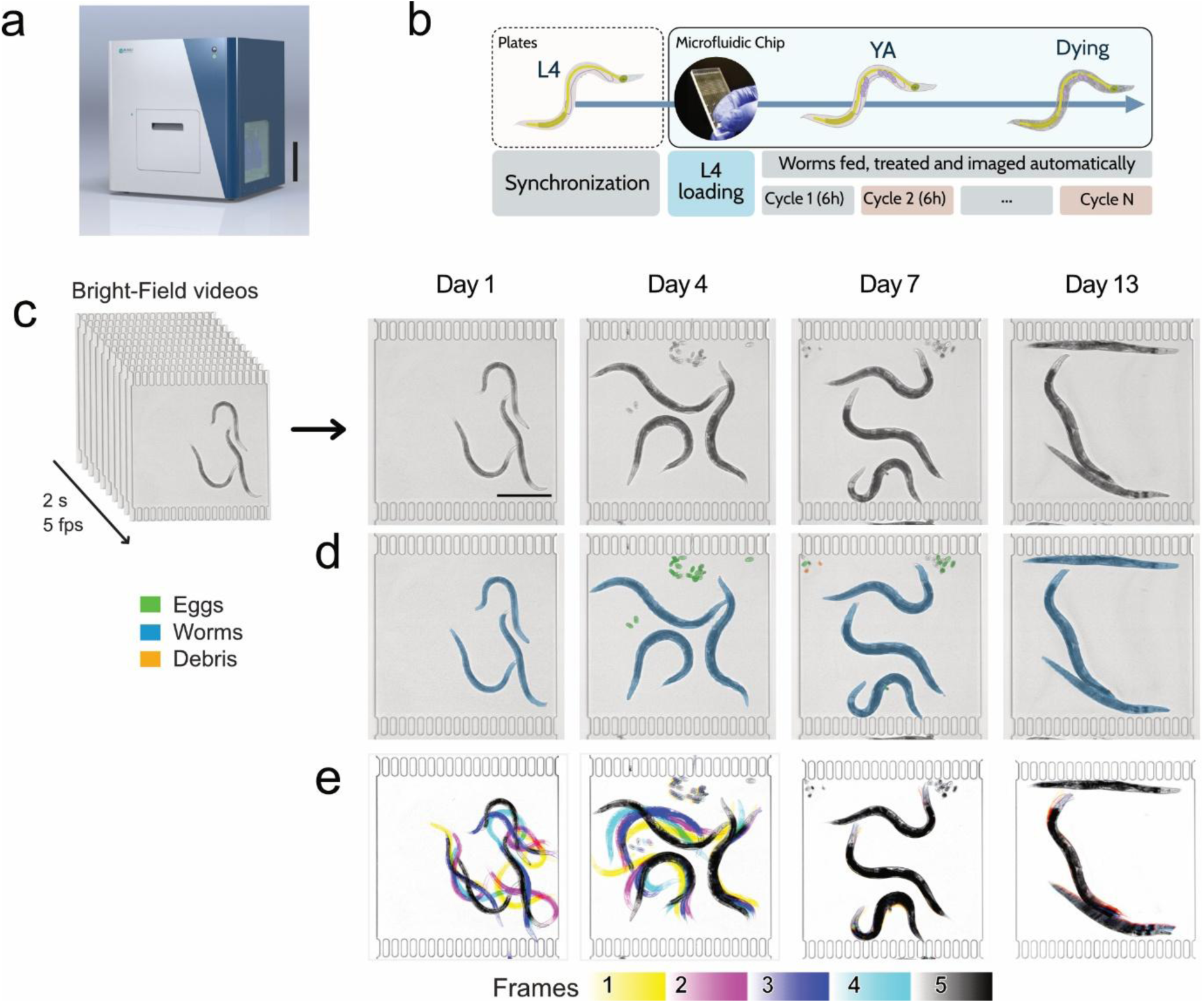
Sydlab^TM^ Device Overview; Growth and reproduction of *C. elegans* N2. **a**,3D rendering of a SydLab^TM^ device. Scale bar, 20cm. **b**, Experimental workflow. YA: young adult. **c**, Brightfield videos acquired every 6 h (2-seconds movie at 5 frames per second) for each microchamber. Example frames were acquired at different time points (Day 1-13, as indicated) from one chamber containing N2 worms. **d**, Computer vision algorithms were applied to single frames to detect different classes of objects (worms, eggs,), with detection masks shown for each class. Note the absence of larvae, developed from viable eggs (green masks), which were eliminated at the L1 stage thanks to the liquid flow. **e**, Five-frame projections (one color per frame) illustrate N2 worm motility within the chambers; five successive frames are shown from a ten-frame recording.

**Fig. S2.**
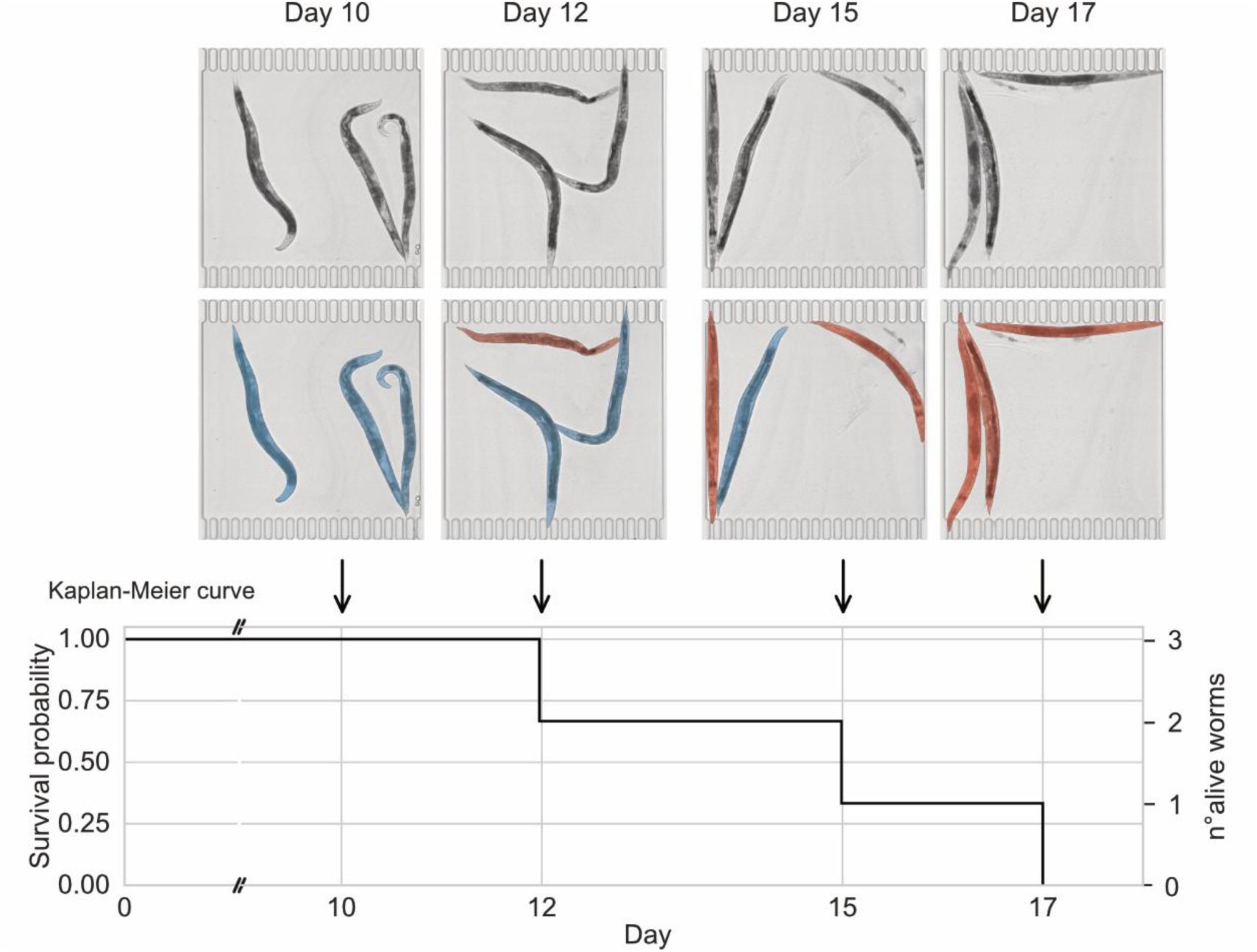
Death detection algorithm. Bright field images at indicated time points showing the same 3 worms, initially alive (blue) and then dead (red), as highlighted with colors on the bottom pictures. The corresponding Kaplan-Meier curves for this chamber is shown.

**Fig. S3.**
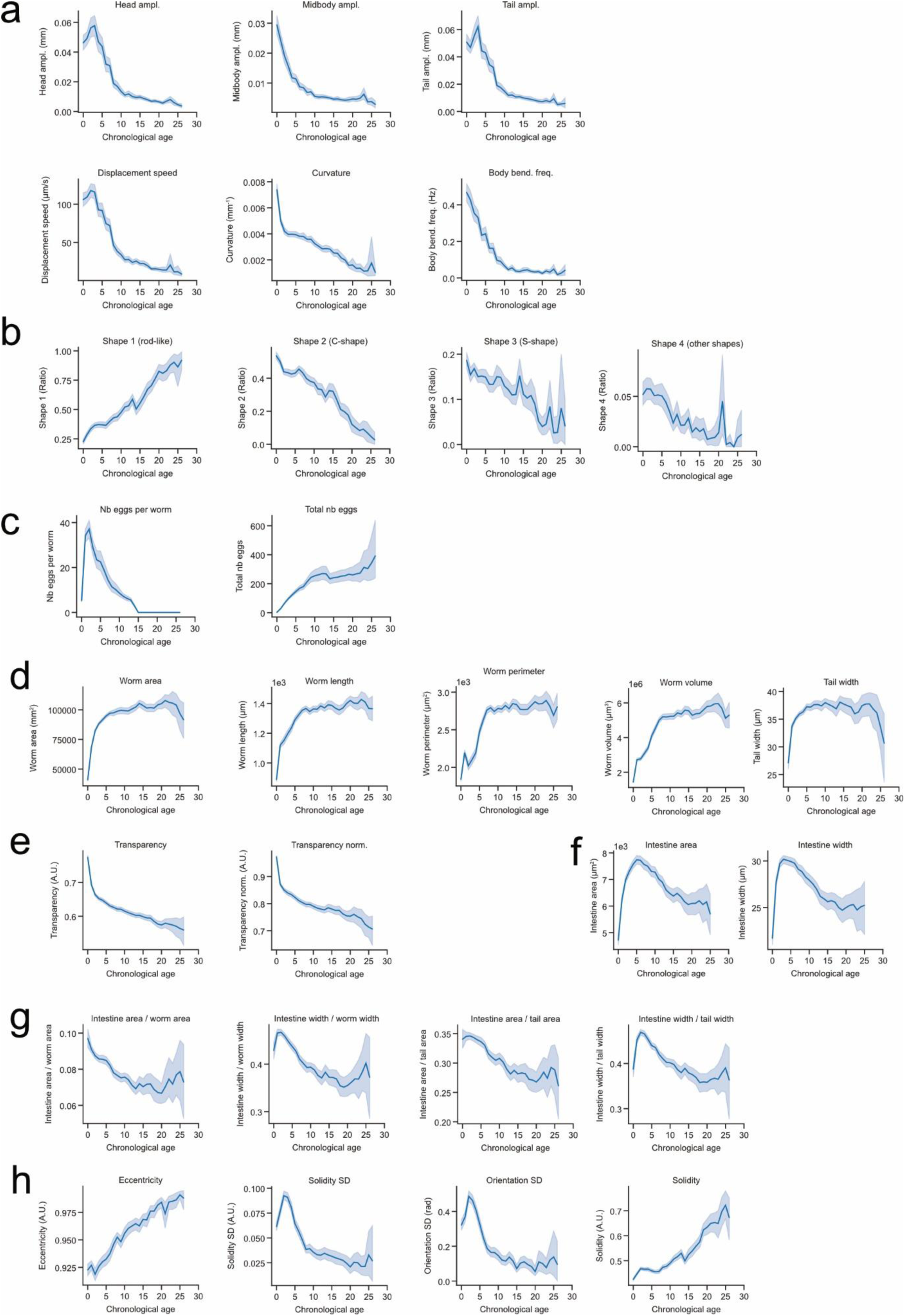
Changes in features with chronological age in control worms. **a-c**, Measurements of selected phenotypic features as a function of time (days) in N2 worms (N=366). Lines represent the mean, shaded areas indicate the 95% confidence interval (CI) of the mean. **a**, motility-related features, **b**, posture, **c**, reproduction, **d**, growth, **e**, transparency, **f**, intestine size, **g**, intestinal features normalized on worm size, **h**, low-level features (LLFs).

**Fig. S4.**
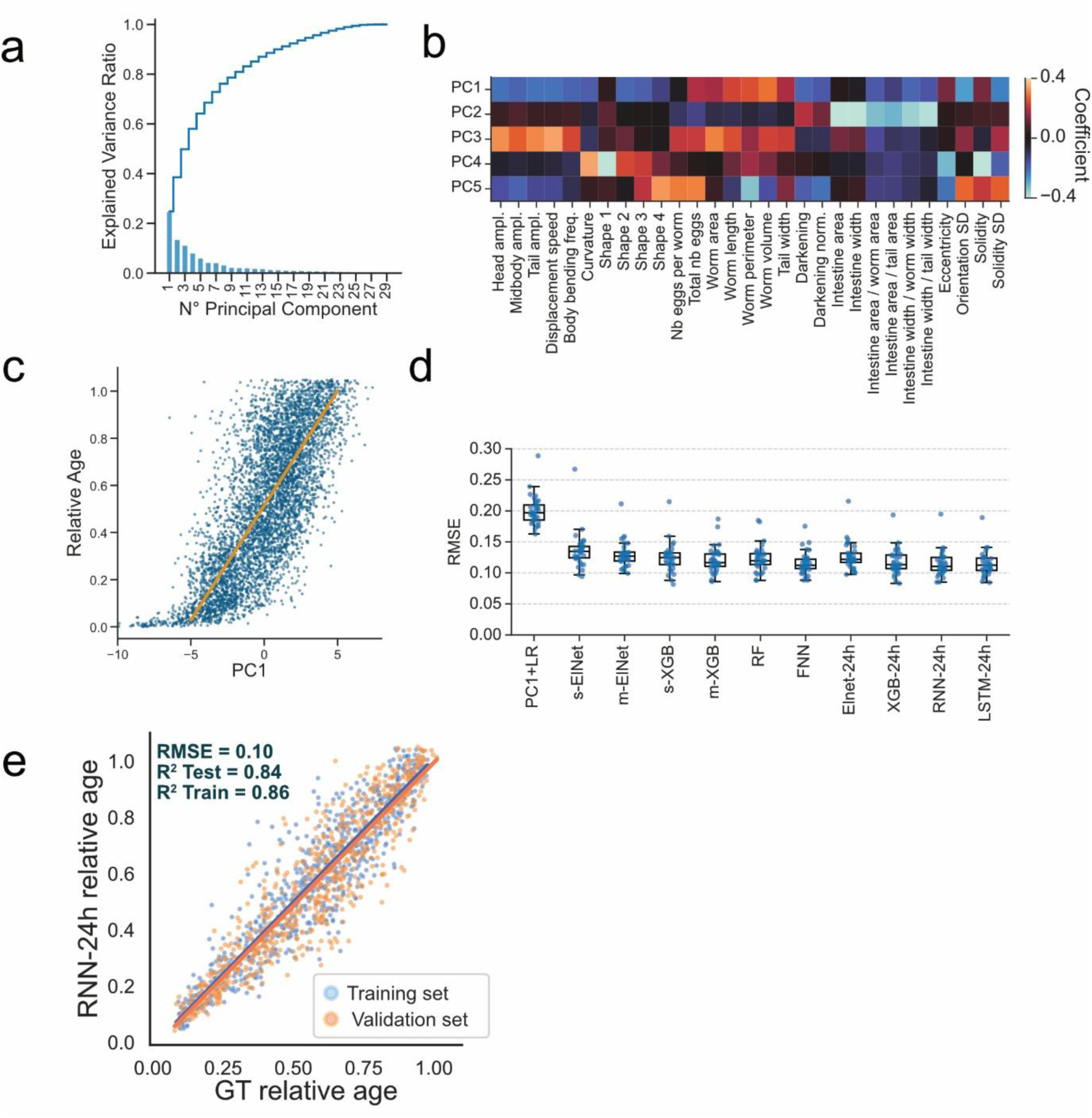
Additional data for PCA, model performances and feature weights. **a**, Explained variance of the first 29 principal components of the PCA of Fig 2a, b. Bars represent the explained variance of one component, the line shows the cumulative explained variance by the successive components. **b**, Heatmap of the loadings per features for the first five principal components of the PCA shown in Fig. 2 a. Color scale indicates loading coefficients. PC2 primarily reflects variation in intestinal traits. **c**, Relative age as a function of PC1. Individual recordings are shown as dots, the result of a least-square linear regression is shown in orange. 5000 points were sampled for graphical clarity. **d**, Root Mean Square Error (RMSE) values obtained from the different models tested. Models were evaluated with LOGOCV (see Methods). **e**, Correlation plot of the RNN-24h model trained on 95% of the dataset (4275 individuals, blue) and tested on the remaining 5% (orange). Each point represents the ground truth (GT) relative age versus the model’s predicted relative age. A least-squares regression (plain line) was fitted to each dataset. For visual clarity, 400 predictions from each set were randomly sampled.

**Fig. S5.**
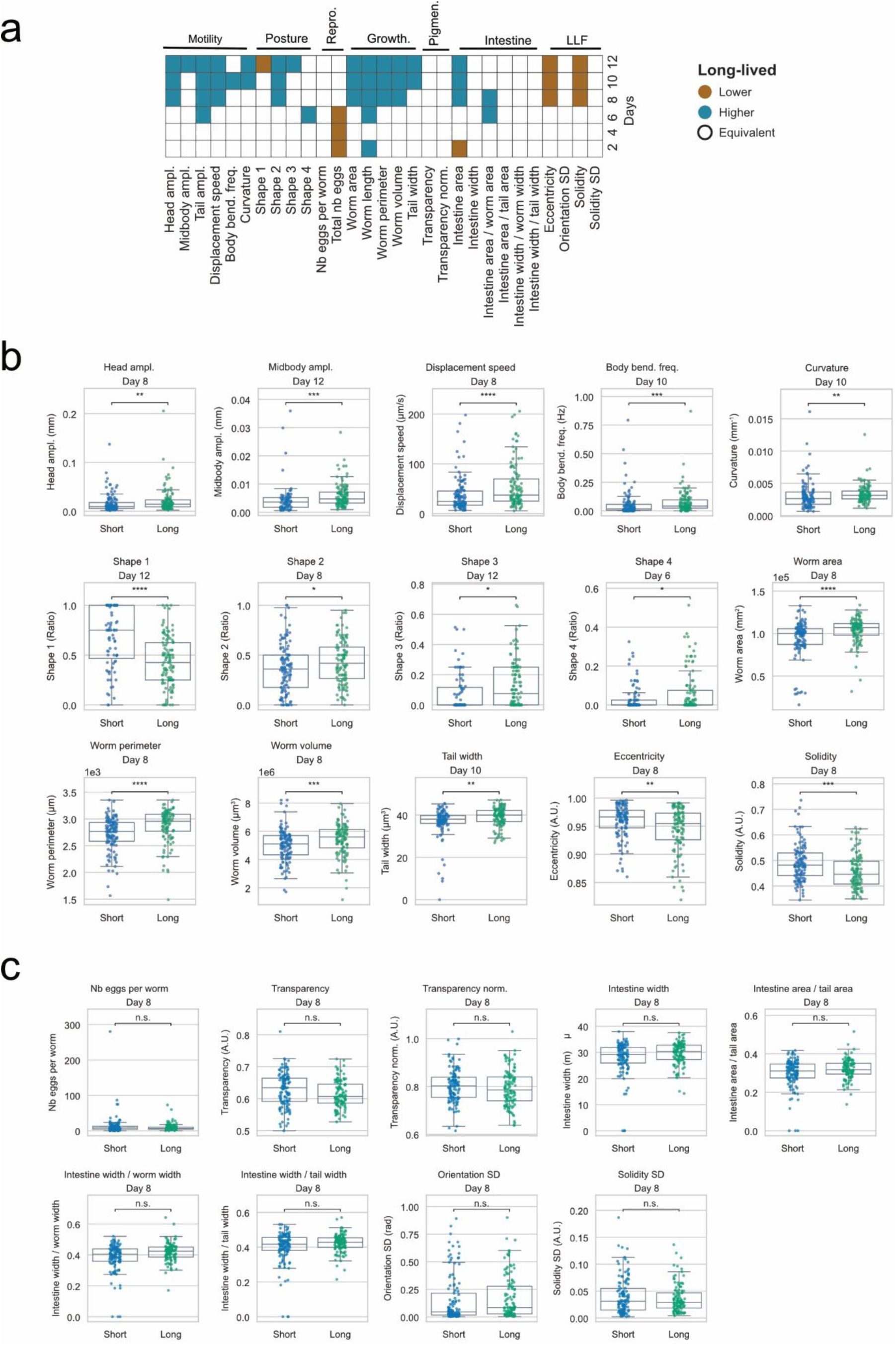
Comparison of features across N2 short and long-lived subpopulations over chronological age. **a**, Heatmap summarizing differences in phenotypic features values between long-lived and short-lived worms across adulthood. For each feature, comparisons between cohorts were performed every two days using two-sided Mann-Whitney U tests, followed by Benjamini-Hochberg correction for multiple testing (feature-wise). At time points showing a statistically significant difference (adjusted p < 0.05), cells are colored according to the direction of the effect: blue indicates higher feature values in long-lived worms, and brown indicates lower values in long-lived worms. Cells are left blank when no significant difference was detected. **b**, boxplots of features with statistically significant difference later in life between short- and long-lived worms. **c**, boxplots at day 8 of features without statistically significant difference detected in the first 12 days between short- and long-lived worms.

**Fig. S6.**
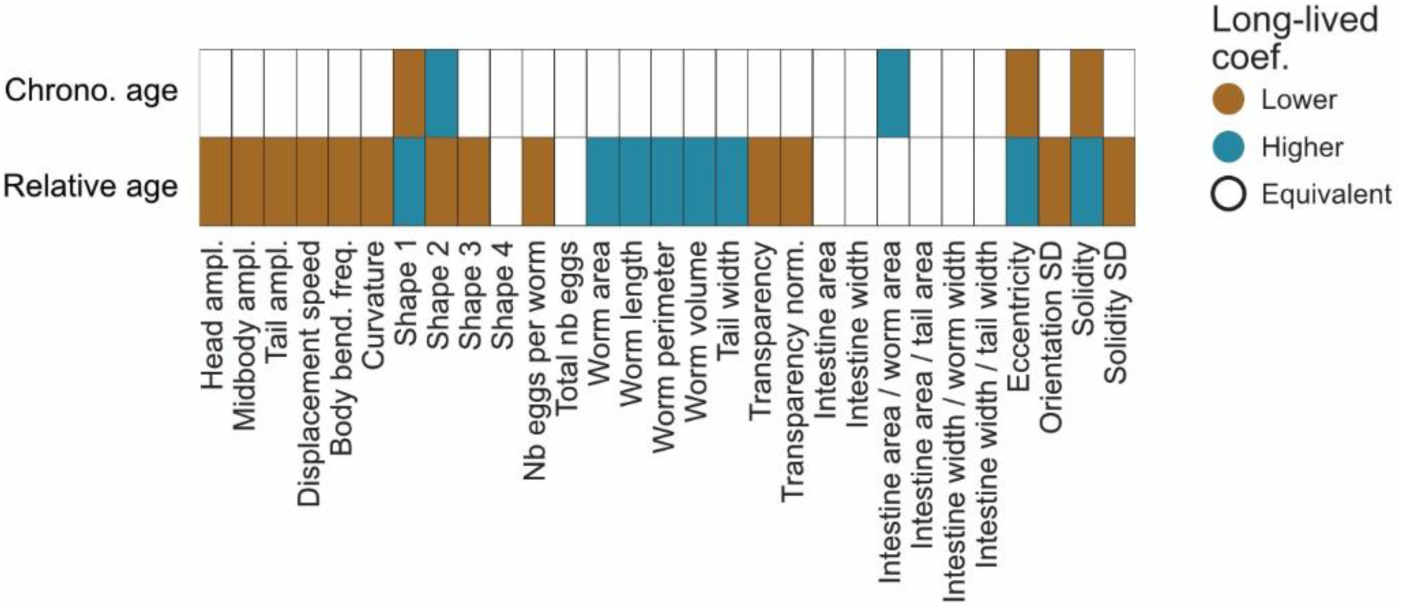
Comparison of features across N2 short and long-lived subpopulations over chronological and relative age using GAMM. Heatmap summarizing differences in phenotypic features between long-lived and short-lived. For each feature, comparisons between cohorts were performed using GAMM (see Methods). Blue indicates significantly higher GAMM coefficient in long-lived worms, and brown indicates significantly lower values.

**Fig. S7.**
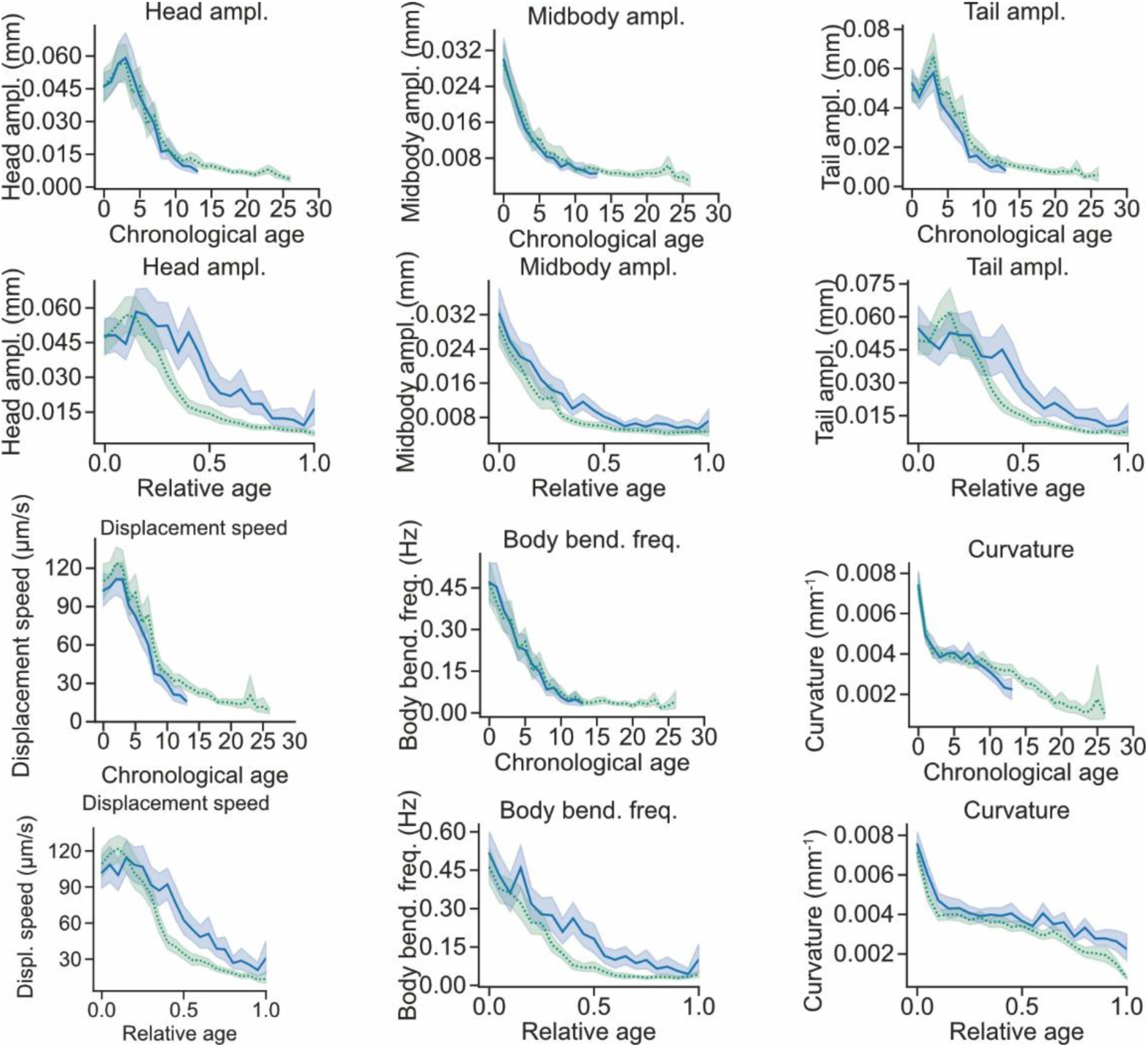
Comparison of features across N2 short and long-lived subpopulations – motility. Age-dependent trajectories of motility features in short- and long-lived worms across adulthood, plotted against chronological age and relative age. Lines indicate the mean and shaded areas the 95% confidence interval of the mean. The same display convention is used in the following plots.

**Fig. S8.**
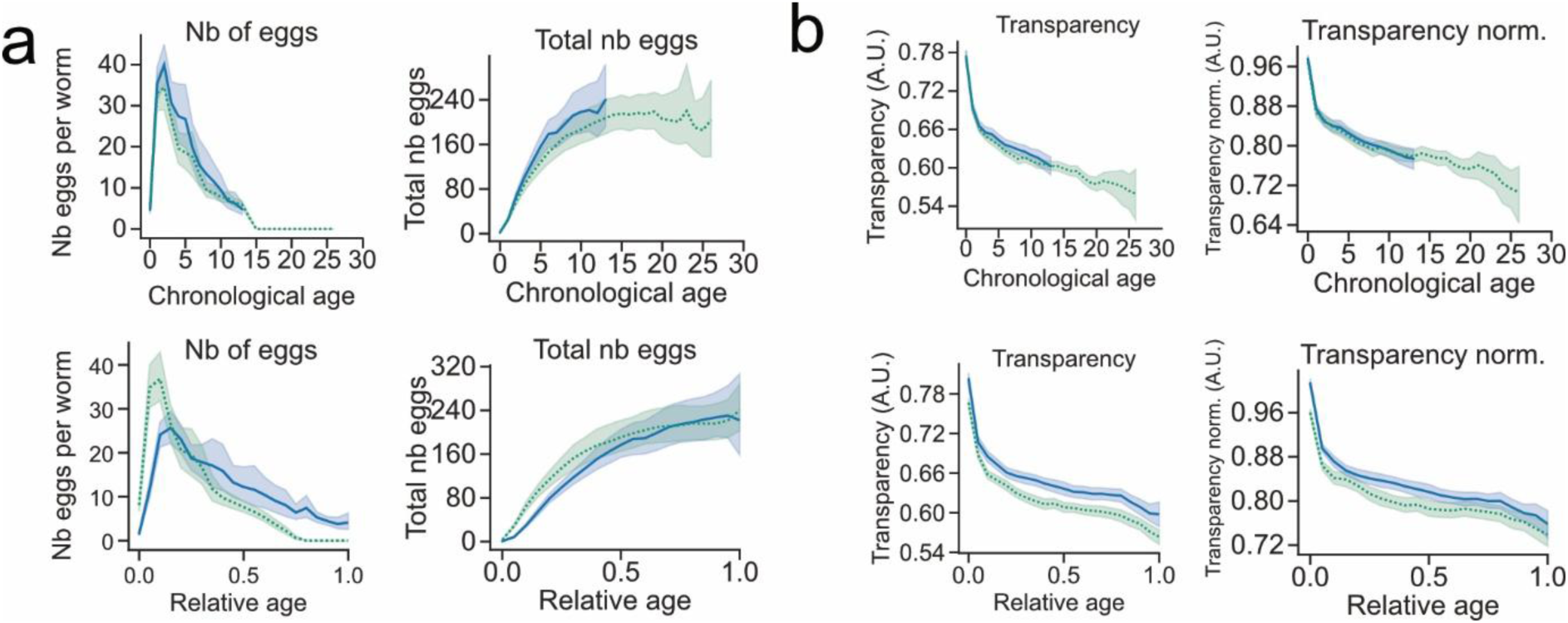
Comparison of features across N2 short and long-lived subpopulations – reproduction and transparency. **a**, reproduction, **b**, transparency

**Fig. S9.**
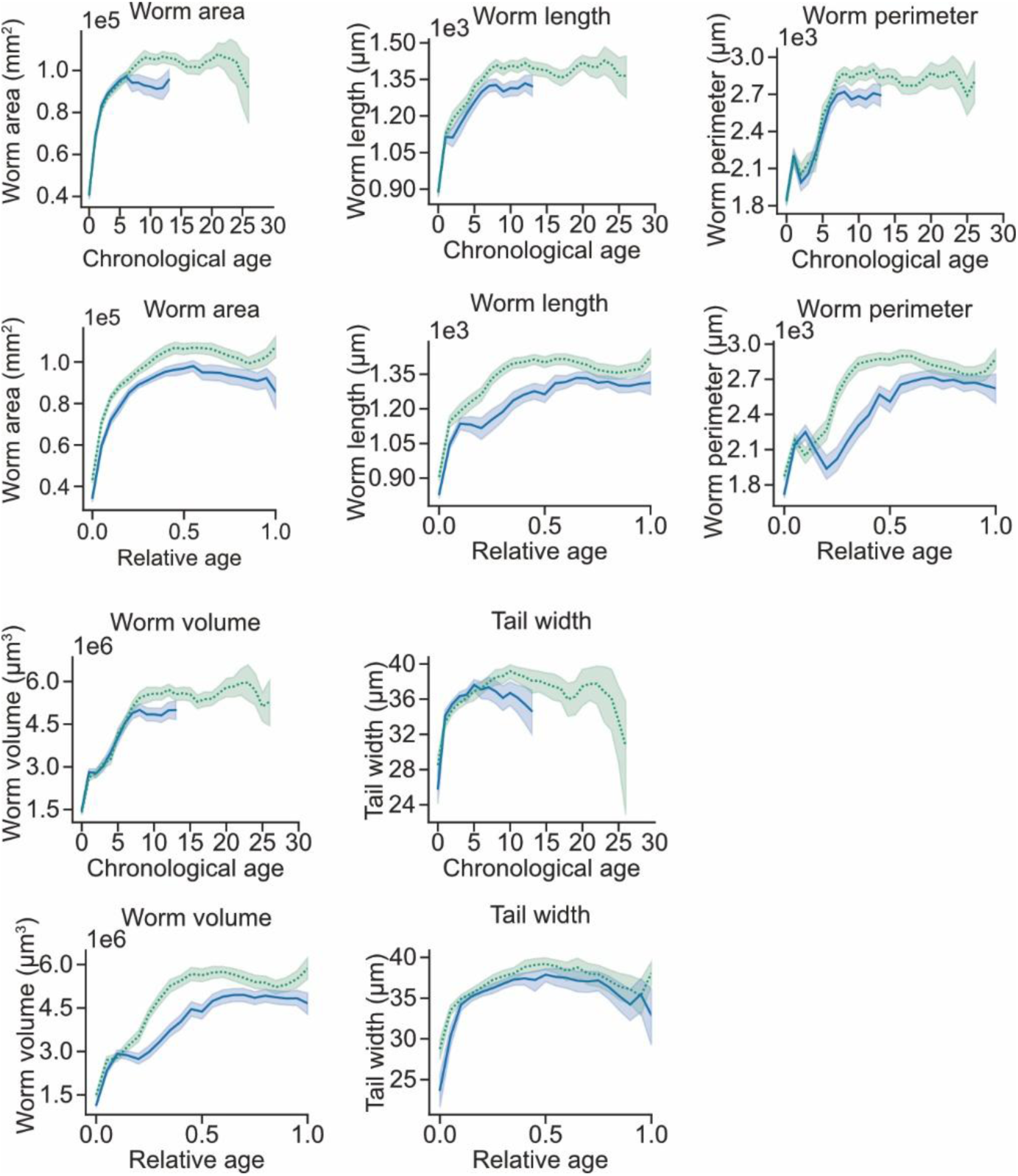
Comparison of features across N2 short and long-lived subpopulations – Growth.

**Fig. S10.**
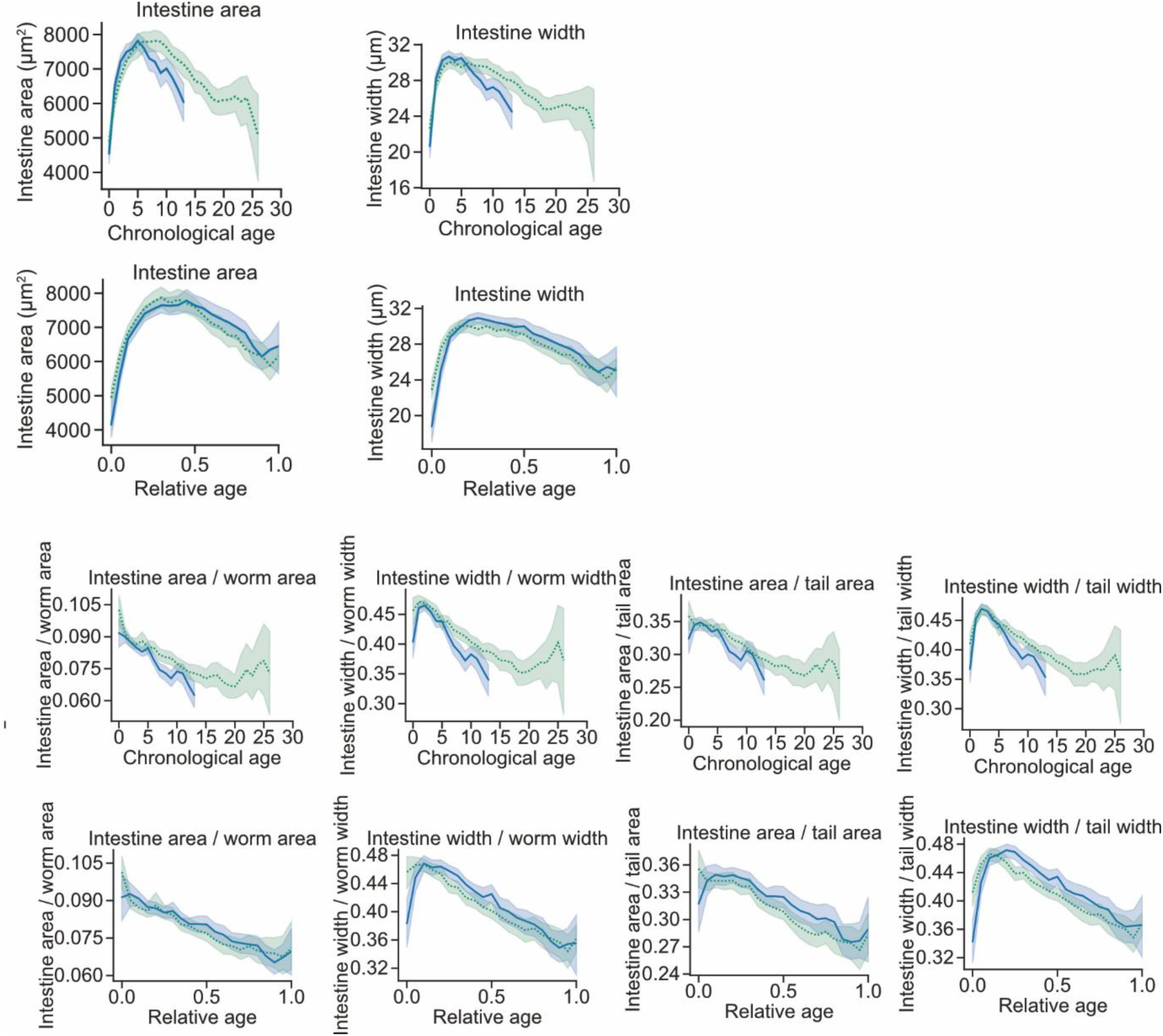
Comparison of features across N2 short and long-lived subpopulations – Intestine.

**Fig. S11.**
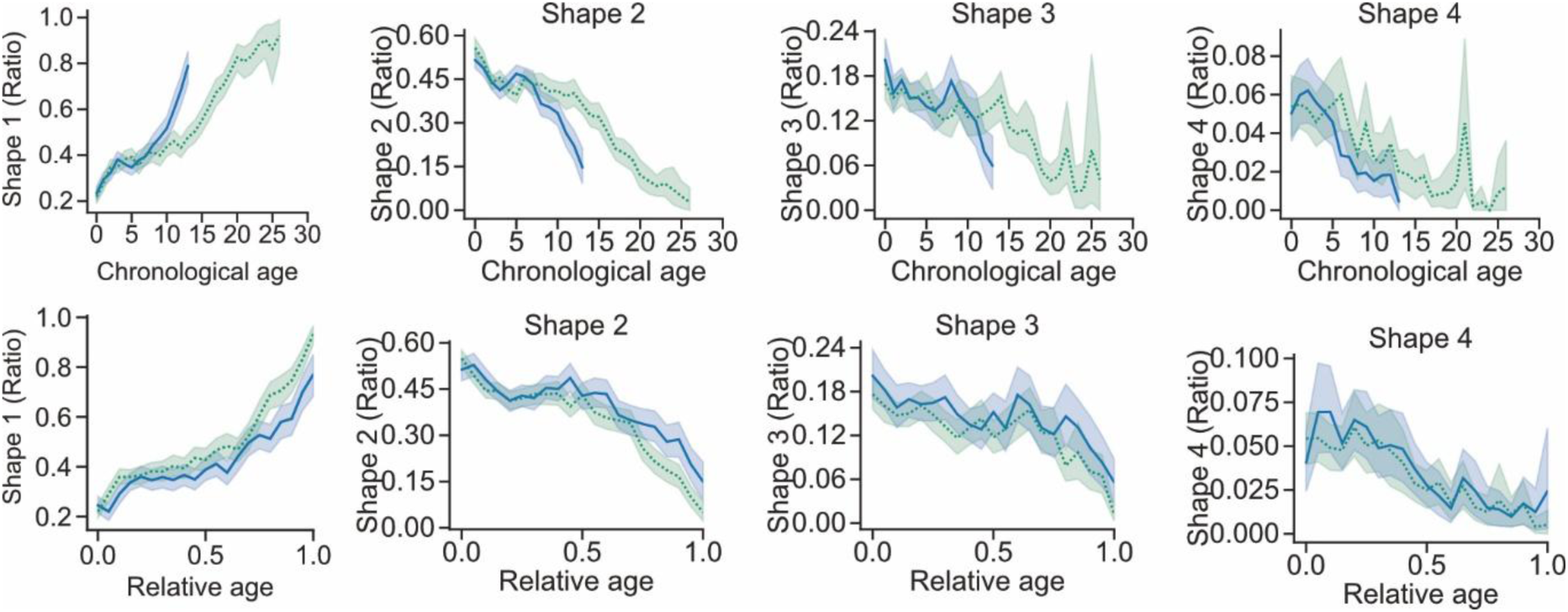
Comparison of features across N2 short and long-lived subpopulations – Posture.

**Fig. S12.**
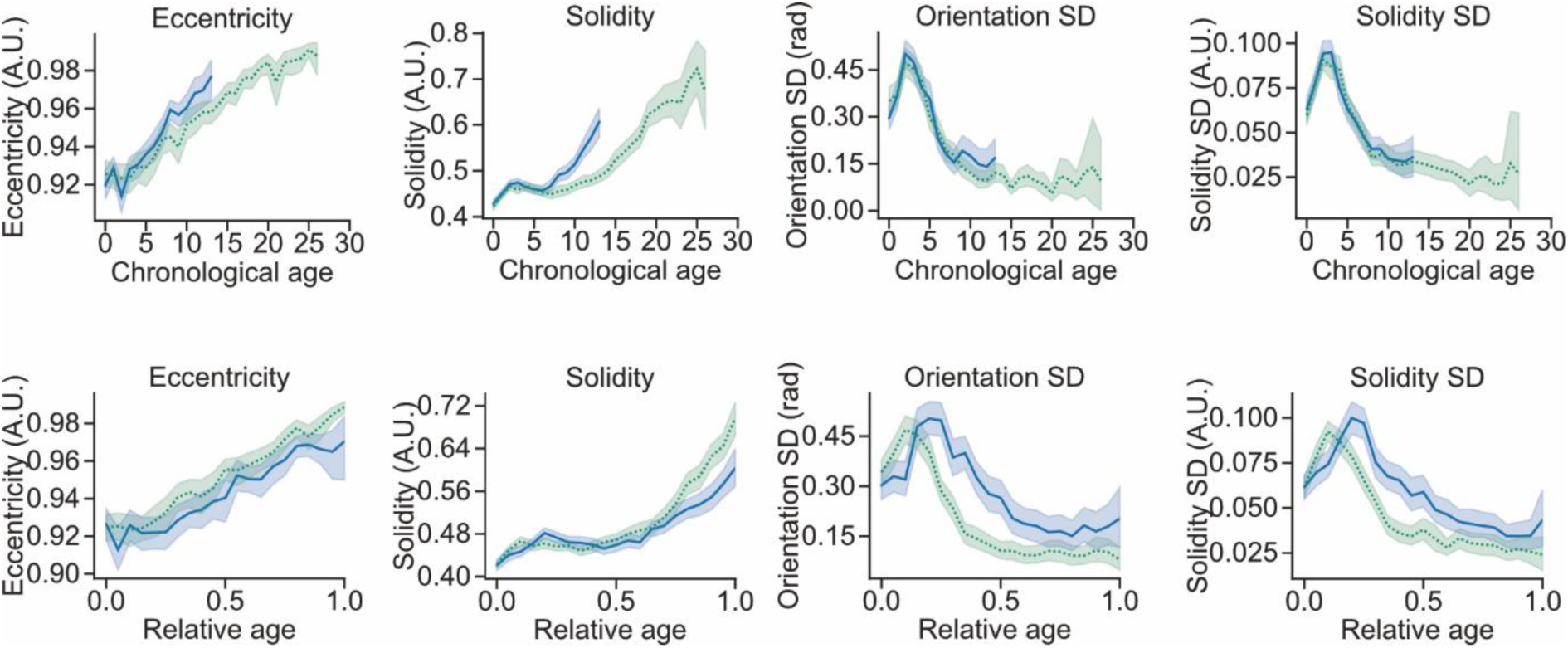
Comparison of features across N2 short and long-lived subpopulations – LLFs.

**Fig. S13.**
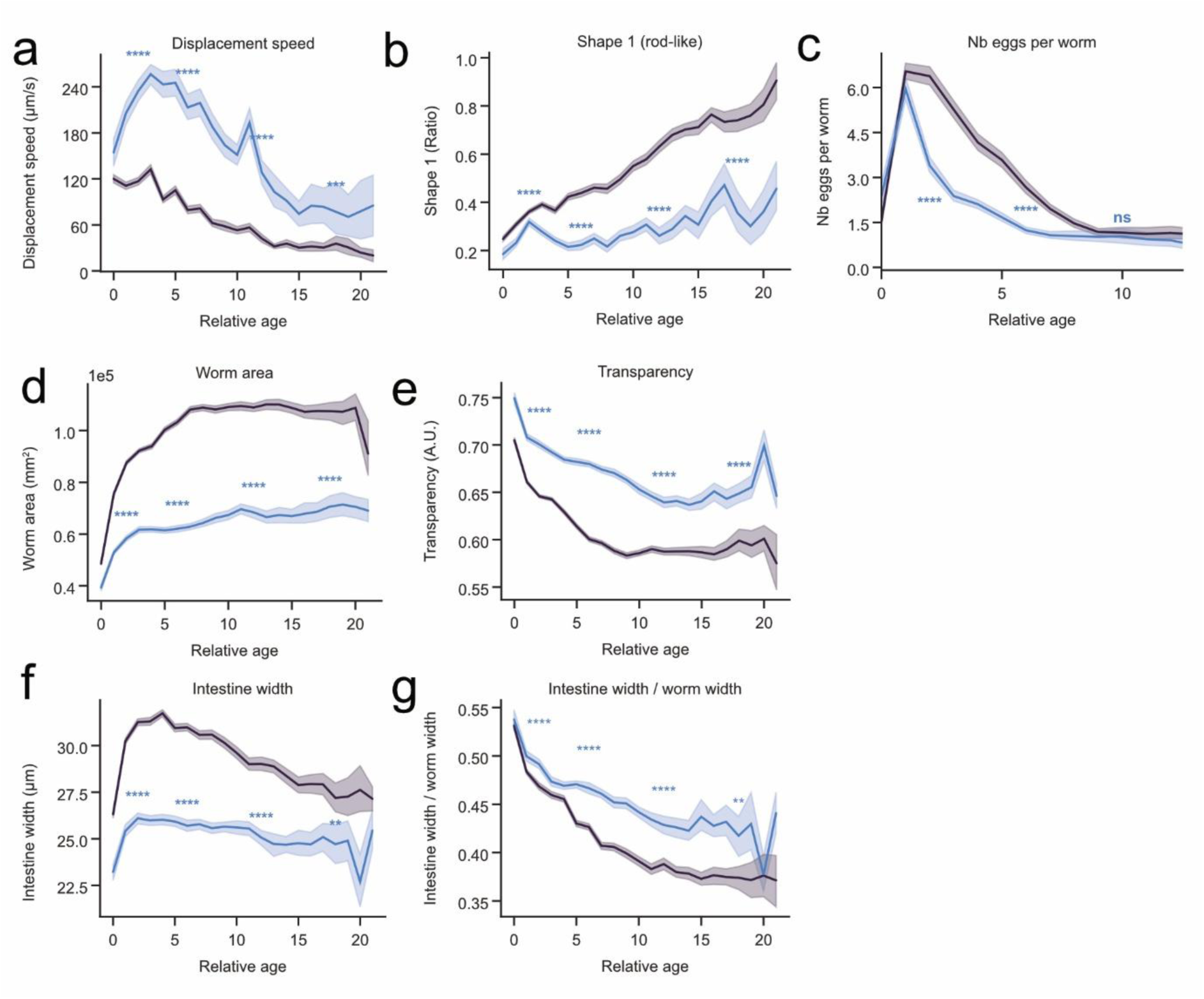
Features dynamics for DR worms. **a**, Displacement speed, **b**, Shape 1 (rod-like), **c**, number of eggs, **d**, worm area, **e**, transparency, **f**, intestine width, **g**, intestine width / worm width Mann–Whitney U tests comparing DR conditions to the AL control at days 2, 6, 12 and 18 (except for eggs where comparisons are done at days 2, 6, and 10). *P* values were corrected for multiple comparisons using the Benjamini-Yekutieli procedure.

**Fig. S14.**
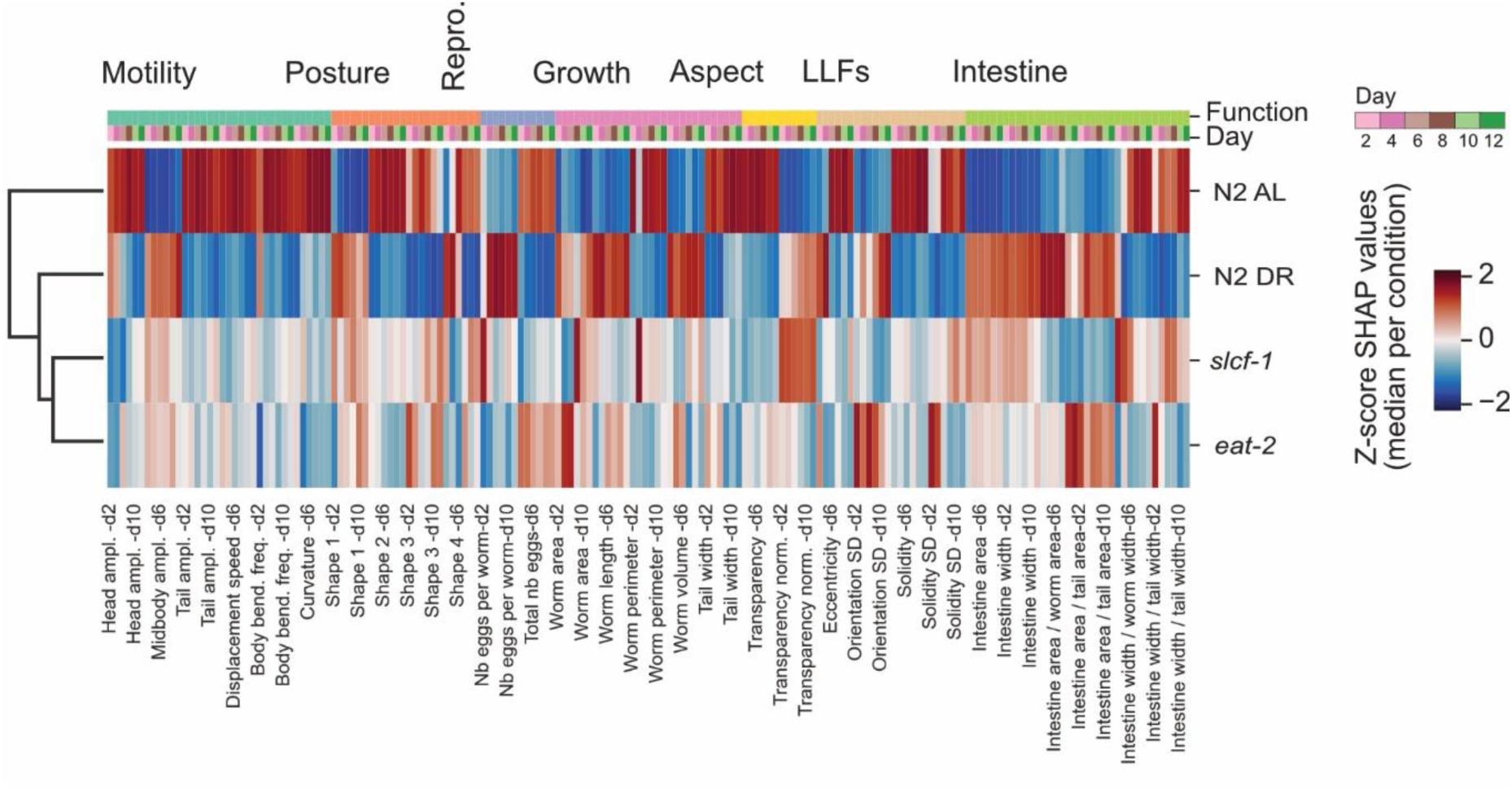
Hierarchical clustering for DR conditions. Heatmap of SHAP using features from days 2, 4, 6, 8, 10, 12. The heatmap shows the median Z-score of features SHAP values for each condition. The dendrogram from hierarchical clustering performed on conditions is shown. Above the heatmap, the trait to which a feature belongs is shown, as well as the time when the feature was recorded.

**Fig. S15.**
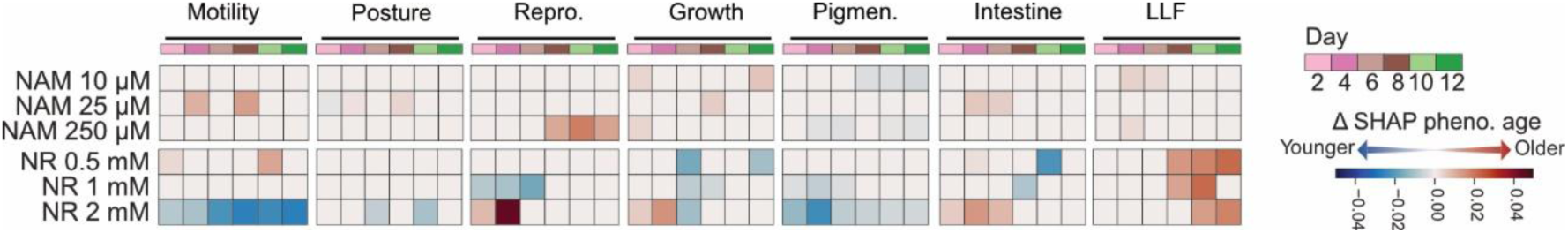
SHAP values for NAD^+^-Boosters. Heatmap of differences in SHAP traits values with controls of NAM at 10, 25, and 250 µM, and NR at 0.5, 1, and 2 mM. Non-significant differences (two-sided Mann-Whitney U test) were set to 0 in the heatmap.

**Tables S1.** Metrics of model performance

|  |  |  |  |  |  |  |  |
| --- | --- | --- | --- | --- | --- | --- | --- |
| Tables S1 Metrics of model performance |  |  |  |  |  |  |  |
| Legend: all = performance on set on worms from all microchamber types |  |  |  |  |  |  |  |
| single = perf. on set with worms alone in their microchambers |  |  |  |  |  |  |  |
| multi = perf. on set with 2 to 3 worms in their microchambers |  |  |  |  |  |  |  |
|  | **R<sup>2</sup> training all** |  |  |  |  |  |  |
| **Model** | **mean** | **std** | **min** | **25%** | **50%** | **75%** | **max** |
| **PC1+LR** | #N/A | #N/A | #N/A | #N/A | #N/A | #N/A | #N/A |
| **s-EINet** | 0.799523 | 0.003235 | 0.794824 | 0.797869 | 0.799711 | 0.800609 | 0.813689 |
| **m-EINet** | 0.811227 | 0.002249 | 0.807268 | 0.810321 | 0.811138 | 0.812105 | 0.82033 |
| **s-XGB** | 0.892636 | 0.001238 | 0.890224 | 0.892112 | 0.89256 | 0.893181 | 0.896814 |
| **m-XGB** | 0.891353 | 0.000956 | 0.889565 | 0.890943 | 0.891306 | 0.891684 | 0.893624 |
| **RF** | 0.918207 | 0.000882 | 0.916604 | 0.917744 | 0.918101 | 0.918636 | 0.920459 |
| **FNN** | 0.858063 | 0.009244 | 0.836589 | 0.852945 | 0.860213 | 0.864153 | 0.875866 |
| **Elnet-24h** | 0.804005 | 0.002485 | 0.79843 | 0.80305 | 0.803694 | 0.805067 | 0.81363 |
| **XGB-24h** | 0.903614 | 0.000868 | 0.901925 | 0.903153 | 0.903563 | 0.903984 | 0.905704 |
| **RNN-24h** | 0.858672 | 0.019591 | 0.817266 | 0.848443 | 0.859436 | 0.871065 | 0.892527 |
| **LSTM-24h** | 0.879787 | 0.027643 | 0.828 | 0.857019 | 0.881072 | 0.905437 | 0.927969 |
|  | **R<sup>2</sup> test all** |  |  |  |  |  |  |
| **Model** | **mean** | **std** | **min** | **25%** | **50%** | **75%** | **max** |
| **PC1+LR** | 0.509324 | 0.136607 | -0.00752 | 0.467433 | 0.534803 | 0.600957 | 0.707414 |
| **s-EINet** | 0.763528 | 0.125525 | 0.146534 | 0.74978 | 0.79075 | 0.812487 | 0.900423 |
| **m-EINet** | 0.789592 | 0.073806 | 0.481758 | 0.750483 | 0.803272 | 0.830308 | 0.883054 |
| **s-XGB** | 0.803239 | 0.084023 | 0.448943 | 0.787718 | 0.813889 | 0.843791 | 0.925125 |
| **m-XGB** | 0.815082 | 0.068147 | 0.609862 | 0.786106 | 0.836154 | 0.856463 | 0.914122 |
| **RF** | 0.808536 | 0.068556 | 0.613582 | 0.782883 | 0.822833 | 0.848701 | 0.915305 |
| **FNN** | 0.830331 | 0.057381 | 0.643668 | 0.814793 | 0.848645 | 0.864827 | 0.914852 |
| **Elnet-24h** | 0.778488 | 0.082824 | 0.424457 | 0.754161 | 0.797779 | 0.822113 | 0.88009 |
| **XGB-24h** | 0.806698 | 0.073126 | 0.536504 | 0.763218 | 0.828971 | 0.848574 | 0.911514 |
| **RNN-24h** | 0.81991 | 0.069848 | 0.53015 | 0.787202 | 0.838215 | 0.858135 | 0.913795 |
| **LSTM-24h** | 0.819322 | 0.067317 | 0.55659 | 0.791259 | 0.838341 | 0.85859 | 0.911897 |
|  | **R<sup>2</sup> test single** |  |  |  |  |  |  |
| **Model** | **mean** | **std** | **min** | **25%** | **50%** | **75%** | **max** |
| **PC1+LR** | 0.509324 | 0.136607 | -0.00752 | 0.467433 | 0.534803 | 0.600957 | 0.707414 |
| **s-EINet** | 0.763528 | 0.125525 | 0.146534 | 0.74978 | 0.79075 | 0.812487 | 0.900423 |
| **m-EINet** | 0.768096 | 0.121018 | 0.163945 | 0.763037 | 0.790405 | 0.817355 | 0.894735 |
| **s-XGB** | 0.803239 | 0.084023 | 0.448943 | 0.787718 | 0.813889 | 0.843791 | 0.925125 |
| **m-XGB** | 0.805432 | 0.077657 | 0.48218 | 0.797289 | 0.818741 | 0.846056 | 0.923757 |
| **RF** | 0.79738 | 0.083933 | 0.429412 | 0.786983 | 0.810426 | 0.83694 | 0.923502 |
| **FNN** | 0.81324 | 0.076602 | 0.487391 | 0.806603 | 0.833636 | 0.851168 | 0.920294 |
| **Elnet-24h** | 0.753765 | 0.135729 | 0.069157 | 0.751509 | 0.777357 | 0.810751 | 0.890894 |
| **XGB-24h** | 0.789317 | 0.103686 | 0.307615 | 0.768017 | 0.811601 | 0.833694 | 0.918482 |
| **RNN-24h** | 0.796961 | 0.099838 | 0.308262 | 0.790788 | 0.815806 | 0.834176 | 0.918092 |
| **LSTM-24h** | 0.797842 | 0.094047 | 0.359815 | 0.781198 | 0.820767 | 0.834723 | 0.913688 |
|  | R^2 test multi |  |  |  |  |  |  |
| Model | mean | std | min | 25% | 50% | 75% | max |
| PC1+LR | #N/A | #N/A | #N/A | #N/A | #N/A | #N/A | #N/A |
| s-EINet | #N/A | #N/A | #N/A | #N/A | #N/A | #N/A | #N/A |
| m-EINet | 0.810348 | 0.060536 | 0.659674 | 0.786971 | 0.823458 | 0.852221 | 0.922071 |
| s-XGB | #N/A | #N/A | #N/A | #N/A | #N/A | #N/A | #N/A |
| m-XGB | 0.822974 | 0.089129 | 0.455768 | 0.784745 | 0.84954 | 0.872795 | 0.930277 |
| RF | 0.818937 | 0.082244 | 0.516469 | 0.785287 | 0.840497 | 0.871375 | 0.940645 |
| FNN | 0.846871 | 0.056933 | 0.70888 | 0.816139 | 0.862987 | 0.886429 | 0.935005 |
| Elnet-24h | 0.802528 | 0.06982 | 0.642533 | 0.781024 | 0.81553 | 0.84245 | 0.93571 |
| XGB-24h | 0.823373 | 0.067709 | 0.693904 | 0.771846 | 0.84466 | 0.865687 | 0.927163 |
| RNN-24h | 0.840388 | 0.063526 | 0.671866 | 0.793659 | 0.856228 | 0.886374 | 0.931508 |
| LSTM-24h | 0.839162 | 0.061308 | 0.701232 | 0.792523 | 0.856178 | 0.88856 | 0.935037 |
|  | RMSE all |  |  |  |  |  |  |
| Model | mean | std | min | 25% | 50% | 75% | max |
| PC1+LR | 0.20055 | 0.023345 | 0.162794 | 0.185333 | 0.196977 | 0.209553 | 0.288866 |
| s-EINet | 0.135587 | 0.02912 | 0.094119 | 0.124159 | 0.134707 | 0.142663 | 0.26722 |
| m-EINet | 0.129412 | 0.021001 | 0.099205 | 0.119245 | 0.126832 | 0.133466 | 0.211334 |
| s-XGB | 0.124155 | 0.023406 | 0.081614 | 0.113535 | 0.124758 | 0.13225 | 0.214721 |
| m-XGB | 0.120897 | 0.021141 | 0.085979 | 0.110626 | 0.116615 | 0.130393 | 0.186868 |
| RF | 0.123147 | 0.021486 | 0.087906 | 0.11361 | 0.119746 | 0.131046 | 0.18454 |
| FNN | 0.116054 | 0.018117 | 0.088123 | 0.107064 | 0.112206 | 0.122089 | 0.175239 |
| Elnet-24h | 0.126951 | 0.022112 | 0.097513 | 0.117004 | 0.12176 | 0.131793 | 0.215578 |
| XGB-24h | 0.118241 | 0.0208 | 0.08302 | 0.106922 | 0.113522 | 0.128935 | 0.193459 |
| RNN-24h | 0.114233 | 0.020331 | 0.085034 | 0.104298 | 0.110568 | 0.124987 | 0.19478 |
| LSTM-24h | 0.114448 | 0.019667 | 0.084542 | 0.104051 | 0.112378 | 0.123785 | 0.189221 |
|  | RMSE test single |  |  |  |  |  |  |
| Model | mean | std | min | 25% | 50% | 75% | max |
| PC1+LR | 0.20055 | 0.023345 | 0.162794 | 0.185333 | 0.196977 | 0.209553 | 0.288866 |
| s-EINet | 0.135587 | 0.02912 | 0.094119 | 0.124159 | 0.134707 | 0.142663 | 0.26722 |
| m-EINet | 0.134528 | 0.028634 | 0.096914 | 0.123246 | 0.132606 | 0.14188 | 0.264212 |
| s-XGB | 0.124155 | 0.023406 | 0.081614 | 0.113535 | 0.124758 | 0.13225 | 0.214721 |
| m-XGB | 0.123682 | 0.022165 | 0.08248 | 0.11371 | 0.123011 | 0.129584 | 0.207934 |
| RF | 0.126234 | 0.023722 | 0.082617 | 0.118347 | 0.127689 | 0.133717 | 0.218271 |
| FNN | 0.121158 | 0.02189 | 0.084332 | 0.112997 | 0.119144 | 0.125079 | 0.206885 |
| Elnet-24h | 0.132486 | 0.029901 | 0.095028 | 0.12172 | 0.130389 | 0.139098 | 0.272108 |
| XGB-24h | 0.122556 | 0.02567 | 0.08214 | 0.109746 | 0.121466 | 0.128043 | 0.23468 |
| RNN-24h | 0.120543 | 0.024884 | 0.082337 | 0.113961 | 0.118388 | 0.12437 | 0.234571 |
| LSTM-24h | 0.120321 | 0.023743 | 0.084521 | 0.112195 | 0.118431 | 0.124273 | 0.22566 |
|  | RMSE test multi |  |  |  |  |  |  |

| Model | mean | std | min | 25% | 50% | 75% | max |
| --- | --- | --- | --- | --- | --- | --- | --- |
| PC1+LR | #N/A | #N/A | #N/A | #N/A | #N/A | #N/A | #N/A |
| s-EINet | #N/A | #N/A | #N/A | #N/A | #N/A | #N/A | #N/A |
| m-EINet | 0.122325 | 0.018866 | 0.078846 | 0.110492 | 0.12061 | 0.131582 | 0.160399 |
| s-XGB | #N/A | #N/A | #N/A | #N/A | #N/A | #N/A | #N/A |
| m-XGB | 0.116737 | 0.026589 | 0.074579 | 0.102253 | 0.112778 | 0.129225 | 0.21928 |
| RF | 0.118353 | 0.025791 | 0.068811 | 0.100573 | 0.114862 | 0.130809 | 0.20669 |
| FNN | 0.109376 | 0.019065 | 0.072006 | 0.094783 | 0.107569 | 0.121539 | 0.151522 |
| Elnet-24h | 0.119125 | 0.020728 | 0.069514 | 0.10616 | 0.116334 | 0.125033 | 0.162979 |
| XGB-24h | 0.112244 | 0.020855 | 0.073991 | 0.100903 | 0.108852 | 0.126273 | 0.157177 |
| RNN-24h | 0.10668 | 0.020154 | 0.07175 | 0.092351 | 0.102371 | 0.116574 | 0.158382 |
| LSTM-24h | 0.107184 | 0.020169 | 0.069877 | 0.093291 | 0.104046 | 0.12194 | 0.151129 |
|  | MAE all |  |  |  |  |  |  |
| Model | mean | std | min | 25% | 50% | 75% | max |
| PC1+LR | 0.160351 | 0.017963 | 0.130375 | 0.150737 | 0.158356 | 0.166896 | 0.230344 |
| s-EINet | 0.083798 | 0.020566 | 0.058945 | 0.071941 | 0.081413 | 0.087281 | 0.174493 |
| m-EINet | 0.079129 | 0.013893 | 0.058249 | 0.06845 | 0.078601 | 0.090009 | 0.121336 |
| s-XGB | 0.071341 | 0.016456 | 0.047294 | 0.060707 | 0.070058 | 0.078541 | 0.133427 |
| m-XGB | 0.06794 | 0.011895 | 0.050662 | 0.058332 | 0.065847 | 0.077198 | 0.094656 |
| RF | 0.06991 | 0.014127 | 0.050412 | 0.056786 | 0.071437 | 0.07982 | 0.106683 |
| FNN | 0.068088 | 0.011956 | 0.05346 | 0.060038 | 0.065378 | 0.076871 | 0.108699 |
| Elnet-24h | 0.080165 | 0.013397 | 0.062926 | 0.070521 | 0.077256 | 0.088815 | 0.120297 |
| XGB-24h | 0.067786 | 0.013343 | 0.048872 | 0.056916 | 0.064064 | 0.079438 | 0.108924 |
| RNN-24h | 0.068895 | 0.014852 | 0.051464 | 0.058771 | 0.063405 | 0.077056 | 0.125904 |
| LSTM-24h | 0.068114 | 0.013821 | 0.051229 | 0.058043 | 0.065503 | 0.078813 | 0.116285 |
|  | MAE test single |  |  |  |  |  |  |
| Model | mean | std | min | 25% | 50% | 75% | max |
| PC1+LR | 0.160351 | 0.017963 | 0.130375 | 0.150737 | 0.158356 | 0.166896 | 0.230344 |
| s-EINet | 0.083798 | 0.020566 | 0.058945 | 0.071941 | 0.081413 | 0.087281 | 0.174493 |
| m-EINet | 0.083051 | 0.019548 | 0.060174 | 0.073146 | 0.078115 | 0.089585 | 0.171323 |
| s-XGB | 0.071341 | 0.016456 | 0.047294 | 0.060707 | 0.070058 | 0.078541 | 0.133427 |
| m-XGB | 0.071701 | 0.015414 | 0.049451 | 0.062044 | 0.070227 | 0.078142 | 0.129306 |
| RF | 0.073732 | 0.017302 | 0.048184 | 0.06162 | 0.074762 | 0.080187 | 0.135403 |
| FNN | 0.072033 | 0.015775 | 0.053006 | 0.062691 | 0.07026 | 0.078524 | 0.141645 |
| Elnet-24h | 0.084141 | 0.019261 | 0.059517 | 0.073222 | 0.079493 | 0.091152 | 0.172581 |
| XGB-24h | 0.071776 | 0.018041 | 0.049222 | 0.060039 | 0.06721 | 0.077925 | 0.145822 |
| RNN-24h | 0.073804 | 0.018464 | 0.050714 | 0.062506 | 0.072945 | 0.081637 | 0.156112 |
| LSTM-24h | 0.072919 | 0.01717 | 0.051767 | 0.061072 | 0.070859 | 0.082405 | 0.142392 |
|  | MAE test multi |  |  |  |  |  |  |
| Model | mean | std | min | 25% | 50% | 75% | max |
| PC1+LR | #N/A | #N/A | #N/A | #N/A | #N/A | #N/A | #N/A |
| s-EINet | #N/A | #N/A | #N/A | #N/A | #N/A | #N/A | #N/A |
| m-EINet | 0.076287 | 0.014628 | 0.043193 | 0.065485 | 0.071667 | 0.089252 | 0.101597 |
| **s-XGB** | #N/A | #N/A | #N/A | #N/A | #N/A | #N/A | #N/A |
| **m-XGB** | 0.06536 | 0.014239 | 0.039793 | 0.056273 | 0.062232 | 0.074834 | 0.116098 |
| **RF** | 0.066917 | 0.016667 | 0.036501 | 0.052965 | 0.066641 | 0.074489 | 0.125521 |
| **FNN** | 0.064326 | 0.011973 | 0.03909 | 0.056997 | 0.062672 | 0.072928 | 0.08805 |
| **Elnet-24h** | 0.077348 | 0.01432 | 0.050054 | 0.06776 | 0.074982 | 0.088429 | 0.105977 |
| **XGB-24h** | 0.064889 | 0.014528 | 0.03763 | 0.055836 | 0.06167 | 0.073486 | 0.108647 |
| **RNN-24h** | 0.064306 | 0.013928 | 0.038337 | 0.056097 | 0.06063 | 0.07502 | 0.104703 |
| **LSTM-24h** | 0.064254 | 0.014561 | 0.04114 | 0.054509 | 0.059942 | 0.074108 | 0.099691 |
|  | **n_worms test set** |  |  |  |  |  |  |
| **Model** | **mean** | **std** | **min** | **25%** | **50%** | **75%** | **max** |
| **PC1+LR** | 62.21212 | 30.21771 | 14 | 40 | 60 | 83 | 140 |
| **s-EINet** | 62.21212 | 30.21771 | 14 | 40 | 60 | 83 | 140 |
| **m-EINet** | 134.9091 | 56.75571 | 37 | 74 | 146 | 164 | 264 |
| **s-XGB** | 62.21212 | 30.21771 | 14 | 40 | 60 | 83 | 140 |
| **m-XGB** | 134.9091 | 56.75571 | 37 | 74 | 146 | 164 | 264 |
| **RF** | 134.9091 | 56.75571 | 37 | 74 | 146 | 164 | 264 |
| **FNN** | 134.9091 | 56.75571 | 37 | 74 | 146 | 164 | 264 |
| **Elnet-24h** | 134.9091 | 56.75571 | 37 | 74 | 146 | 164 | 264 |
| **XGB-24h** | 134.9091 | 56.75571 | 37 | 74 | 146 | 164 | 264 |
| **RNN-24h** | 134.9091 | 56.75571 | 37 | 74 | 146 | 164 | 264 |
| **LSTM-24h** | 134.9091 | 56.75571 | 37 | 74 | 146 | 164 | 264 |
|  | **n_worms multi** |  |  |  |  |  |  |
| **Model** | **mean** | **std** | **min** | **25%** | **50%** | **75%** | **max** |
| **PC1+LR** | 0 | 0 | 0 | 0 | 0 | 0 | 0 |
| **s-EINet** | 0 | 0 | 0 | 0 | 0 | 0 | 0 |
| **m-EINet** | 72.69697 | 38.27326 | 6 | 39 | 69 | 102 | 135 |
| **s-XGB** | 0 | 0 | 0 | 0 | 0 | 0 | 0 |
| **m-XGB** | 72.69697 | 38.27326 | 6 | 39 | 69 | 102 | 135 |
| **RF** | 72.69697 | 38.27326 | 6 | 39 | 69 | 102 | 135 |
| **FNN** | 72.69697 | 38.27326 | 6 | 39 | 69 | 102 | 135 |
| **Elnet-24h** | 72.69697 | 38.27326 | 6 | 39 | 69 | 102 | 135 |
| **XGB-24h** | 72.69697 | 38.27326 | 6 | 39 | 69 | 102 | 135 |
| **RNN-24h** | 72.69697 | 38.27326 | 6 | 39 | 69 | 102 | 135 |
| **LSTM-24h** | 72.69697 | 38.27326 | 6 | 39 | 69 | 102 | 135 |

## References

1. Magalhães, J. P. de, Stevens, M. & Thornton, D. The Business of Anti-Aging Science. Trends Biotechnol. 35, 1062–1073 (2017).

2. Kenyon, C. The first long-lived mutants: discovery of the insulin/IGF-1 pathway for ageing. Philos. Trans. R. Soc. B Biol. Sci. 366, 9–16 (2011).

3. Yang, Y. et al. Metformin decelerates aging clock in male monkeys. Cell 187, 6358–6378.e29 (2024).

4. Fontana, L., Partridge, L. & Longo, V. D. Extending healthy life span--from yeast to humans. Science 328, 321–326 (2010).

5. Yanai, H., Budovsky, A., Barzilay, T., Tacutu, R. & Fraifeld, V. E. Wide-scale comparative analysis of longevity genes and interventions. Aging Cell 16, 1267–1275 (2017).

6. Flachsbart, F. et al. Association of *FOXO3A* variation with human longevity confirmed in German centenarians. Proc. Natl. Acad. Sci. 106, 2700–2705 (2009).

7. Bansal, A., Zhu, L. J., Yen, K. & Tissenbaum, H. A. Uncoupling lifespan and healthspan in Caenorhabditis elegans longevity mutants. Proc. Natl. Acad. Sci. 112, E277–E286 (2015).

8. Banse, S. A. et al. The coupling between healthspan and lifespan in *Caenorhabditis* depends on complex interactions between compound intervention and genetic background. Aging 16, 5829–5855 (2024).

9. Garmany, A., Yamada, S. & Terzic, A. Longevity leap: mind the healthspan gap. Npj Regen. Med. 6, 57 (2021).

10. Garmany, A. & Terzic, A. Healthspan-lifespan gap differs in magnitude and disease contribution across world regions. Commun. Med. 5, 381 (2025).

11. Palmer, R. D. Aging clocks & mortality timers, methylation, glycomic, telomeric and more. A window to measuring biological age. Aging Med. 5, 120–125 (2022).

12. Silva, N. et al. Measuring healthy ageing: current and future tools. Biogerontology 24, 845–866 (2023).

13. Han, J.-D. J. The ticking of aging clocks. Trends Endocrinol. Metab. 35, 11–22 (2024).

14. Meng, D., Zhang, S., Huang, Y., Mao, K. & Han, J.-D. J. Application of AI in biological age prediction. Curr. Opin. Struct. Biol. 85, 102777 (2024).

15. Meyer, D. H., Maklakov, A. A. & Schumacher, B. Aging by the clock and yet without a program. *Nat*. Aging 1–11 (2025) doi:10.1038/s43587-025-00975-2.

16. Li, G. et al. Predicting healthspan and disease risks through biological age. Trends Mol. Med. 0, (2025).

17. Apsley, A. T., Etzel, L., Ye, Q. & Shalev, I. From population science to the clinic? Limits of epigenetic clocks as personal biomarkers. Epigenomics 17, 1447–1461 (2025).

18. Cruz-González, S. et al. Methylation Clocks Do Not Predict Age or Alzheimer’s Disease Risk Across Genetically Admixed Individuals. eLife 14, (2025).

19. Meyer, D. H. & Schumacher, B. BiT age: A transcriptome-based aging clock near the theoretical limit of accuracy. Aging Cell 20, e13320 (2021).

20. Bulteau, R. & Francesconi, M. Real age prediction from the transcriptome with RAPToR. Nat. Methods 19, 969–975 (2022).

21. Gao, S. M. et al. Aging atlas reveals cell-type-specific effects of pro-longevity strategies. *Nat*. Aging 4, 998–1013 (2024).

22. Martineau, C. N., Brown, A. E. X. & Laurent, P. Multidimensional phenotyping predicts lifespan and quantifies health in Caenorhabditis elegans. PLOS Comput. Biol. 16, e1008002 (2020).

23. Dhondt, I. et al. Prediction of biological age by morphological staging of sarcopenia in Caenorhabditis elegans. Dis. Model. Mech. 14, dmm049169 (2021).

24. Morrow, C. S. et al. Endogenous mitochondrial NAD(P)H fluorescence can predict lifespan. *Commun*. Biol. 7, 1–10 (2024).

25. Kern, C. C. et al. Machine learning predicts lifespan and suggests underlying causes of death in aging C. elegans. *Commun*. Biol. 8, 1630 (2025).

26. Yan, C. et al. Fluorescence lifetime clocks quantify senescence and aging. *Nat*. Aging 5, 2532–2545 (2025).

27. Pincus, Z., Smith-Vikos, T. & Slack, F. J. MicroRNA Predictors of Longevity in Caenorhabditis elegans. PLOS Genet. 7, e1002306 (2011).

28. Palikaras, K. et al. Ectopic fat deposition contributes to age-associated pathology in *Caenorhabditis elegans*. J. Lipid Res. 58, 72–80 (2017).

29. Lu, A. T. et al. DNA methylation GrimAge strongly predicts lifespan and healthspan. Aging 11, 303–327 (2019).

30. Chen, T. & Guestrin, C. XGBoost: A Scalable Tree Boosting System. in Proceedings of the 22nd ACM SIGKDD International Conference on Knowledge Discovery and Data Mining 785–794 (2016). doi:10.1145/2939672.2939785.

31. Mienye, I. D., Swart, T. G. & Obaido, G. Recurrent Neural Networks: A Comprehensive Review of Architectures, Variants, and Applications. Information 15, 517 (2024).

32. Scharf, A., Pohl, F., Egan, B. M., Kocsisova, Z. & Kornfeld, K. Reproductive Aging in Caenorhabditis elegans: From Molecules to Ecology. Front. Cell Dev. Biol. 9, (2021).

33. Herndon, L. A. et al. Stochastic and genetic factors influence tissue-specific decline in ageing C. elegans. Nature 419, 808–814 (2002).

34. Churgin, M. A. et al. Longitudinal imaging of Caenorhabditis elegans in a microfabricated device reveals variation in behavioral decline during aging. eLife 6, e26652 (2017).

35. Zhang, W. B. et al. Extended Twilight among Isogenic C. elegans Causes a Disproportionate Scaling between Lifespan and Health. Cell Syst. 3, 333–345.e4 (2016).

36. Stroustrup, N. et al. The temporal scaling of Caenorhabditis elegans ageing. Nature 530, 103–107 (2016).

37. Roth, L. W. & Polotsky, A. J. Can we live longer by eating less? A review of caloric restriction and longevity. Maturitas 71, 315–319 (2012).

38. Cabreiro, F. & Gems, D. Worms need microbes too: microbiota, health and aging in Caenorhabditis elegans. EMBO Mol. Med. 5, 1300–1310 (2013).

39. Klass, M. R. Aging in the nematode Caenorhabditis elegans: Major biological and environmental factors influencing life span. Mech. Ageing Dev. 6, 413–429 (1977).

40. Avery, L. The genetics of feeding in Caenorhabditis elegans. Genetics 133, 897–917 (1993).

41. Mouchiroud, L. et al. Pyruvate imbalance mediates metabolic reprogramming and mimics lifespan extension by dietary restriction in Caenorhabditis elegans. Aging Cell 10, 39–54 (2011).

42. Pontoizeau, C. et al. Metabolomics analysis uncovers that dietary restriction buffers metabolic changes associated with aging in Caenorhabditis elegans. J. Proteome Res. 13, 2910–2919 (2014).

43. Madeo, F., Carmona-Gutierrez, D., Hofer, S. J. & Kroemer, G. Caloric Restriction Mimetics against Age-Associated Disease: Targets, Mechanisms, and Therapeutic Potential. Cell Metab. 29, 592–610 (2019).

44. Pryor, R. et al. Host-Microbe-Drug-Nutrient Screen Identifies Bacterial Effectors of Metformin Therapy. Cell 178, 1299–1312.e29 (2019).

45. Martin-Montalvo, A. et al. Metformin improves healthspan and lifespan in mice. Nat. Commun. 4, 2192 (2013).

46. Fang, E. F. et al. NAD+ in Aging: Molecular Mechanisms and Translational Implications. Trends Mol. Med. 23, 899–916 (2017).

47. Membrez, M. et al. Trigonelline is an NAD+ precursor that improves muscle function during ageing and is reduced in human sarcopenia. Nat. Metab. 1–15 (2024) doi:10.1038/s42255-024-00997-x.

48. San-Miguel, A. & Lu, H. Microfluidics as a tool for C. elegans research. WormBook Online Rev. C Elegans Biol. 1–19 (2013) doi:10.1895/wormbook.1.162.1.

49. Cornaglia, M., Lehnert, T. & Gijs, M. A. M. Microfluidic systems for high-throughput and high-content screening using the nematode Caenorhabditis elegans. Lab. Chip 17, 3736–3759 (2017).

50. Bulterijs, S. & Braeckman, B. P. Phenotypic Screening in C. elegans as a Tool for the Discovery of New Geroprotective Drugs. Pharmaceuticals 13, 164 (2020).

51. Jeon, J., Jang, H., Ryu, H., Jeon, T.-J. & Kim, S. M. Recent Advances in Microfluidic Platforms for C. Elegans Phenotyping: Comprehensive Review of Imaging Technologies and AI-Driven Analysis. BioChip J. 10.1007/s13206-025-00246-7 (2025) doi:10.1007/s13206-025-00246-7.

52. Huang, C., Xiong, C. & Kornfeld, K. Measurements of age-related changes of physiological processes that predict lifespan of Caenorhabditis elegans. Proc. Natl. Acad. Sci. 101, 8084–8089 (2004).

53. Collins, J. J., Huang, C., Hughes, S. & Kornfeld, K. The measurement and analysis of age-related changes in Caenorhabditis elegans. WormBook Online Rev. C Elegans Biol. 1–21 (2008) doi:10.1895/wormbook.1.137.1.

54. Stephens, G. J., Johnson-Kerner, B., Bialek, W. & Ryu, W. S. Dimensionality and Dynamics in the Behavior of C. elegans. PLOS Comput. Biol. 4, e1000028 (2008).

55. Hughes, S. E., Evason, K., Xiong, C. & Kornfeld, K. Genetic and Pharmacological Factors That Influence Reproductive Aging in Nematodes. PLOS Genet. 3, e25 (2007).

56. Garigan, D. et al. Genetic Analysis of Tissue Aging in Caenorhabditis elegans: A Role for Heat-Shock Factor and Bacterial Proliferation. Genetics 161, 1101–1112 (2002).

57. McGee, M. D. et al. Loss of intestinal nuclei and intestinal integrity in aging C. elegans. Aging Cell 10, 699–710 (2011).

58. Ezcurra, M. et al. C. elegans Eats Its Own Intestine to Make Yolk Leading to Multiple Senescent Pathologies. Curr. Biol. 28, 2544–2556.e5 (2018).

59. Zhai, C. et al. Fusion and expansion of vitellogenin vesicles during Caenorhabditis elegans intestinal senescence. Aging Cell 21, e13719 (2022).

60. Kocsisova, Z. et al. How to measure, analyze, and interpret age-related changes in Caenorhabditis elegans: Lessons for mechanistic and evolutionary theories of aging. Mech. Ageing Dev. 229, 112146 (2026).

61. Tarkhov, A. E. et al. A universal transcriptomic signature of age reveals the temporal scaling of Caenorhabditis elegans aging trajectories. Sci. Rep. 9, 7368 (2019).

62. Statzer, C., Reichert, P., Dual, J. & Ewald, C. Y. Longevity interventions temporally scale healthspan in Caenorhabditis elegans. iScience 25, 103983 (2022).

63. Wang, X. et al. Ageing induces tissue-specific transcriptomic changes in Caenorhabditis elegans. EMBO J. 41, EMBJ2021109633 (2022).

64. Yang, Y. et al. Compression of morbidity by interventions that steepen the survival curve. Nat. Commun. 16, 3340 (2025).

65. Coburn, C. et al. Anthranilate Fluorescence Marks a Calcium-Propagated Necrotic Wave That Promotes Organismal Death in C. elegans. PLOS Biol. 11, e1001613 (2013).

66. Galimov, E. R. et al. Coupling of Rigor Mortis and Intestinal Necrosis during C. elegans Organismal Death. Cell Rep. 22, 2730–2741 (2018).

67. Venz, R., Pekec, T., Katic, I., Ciosk, R. & Ewald, C. Y. End-of-life targeted degradation of DAF-2 insulin/IGF-1 receptor promotes longevity free from growth-related pathologies. eLife 10, e71335 (2021).

68. Sultanova, Z. et al. Optimising Age-Specific Insulin Signalling to Slow Down Reproductive Ageing Increases Fitness in Different Nutritional Environments. Aging Cell 24, e14481 (2025).

69. Riddle, D. L., Swanson, M. M. & Albert, P. S. Interacting genes in nematode dauer larva formation. Nature 290, 668–671 (1981).

70. Krajacic, P., Shen, X., Purohit, P. K., Arratia, P. & Lamitina, T. Biomechanical Profiling of Caenorhabditis elegans Motility. Genetics 191, 1015–1021 (2012).

71. Walt, S. van der et al. scikit-image: Image processing in Python. PeerJ 2, e453 (2014).

72. Pedregosa, F. et al. Scikit-learn: Machine Learning in Python. Mach. Learn. PYTHON.

73. Virtanen, P. et al. SciPy 1.0: fundamental algorithms for scientific computing in Python. Nat. Methods 17, 261–272 (2020).

74. Akiba, T., Sano, S., Yanase, T., Ohta, T. & Koyama, M. Optuna: A Next-generation Hyperparameter Optimization Framework. in Proceedings of the 25th ACM SIGKDD International Conference on Knowledge Discovery & Data Mining 2623–2631 (Association for Computing Machinery, New York, NY, USA, 2019). doi:10.1145/3292500.3330701.

75. Paszke, A., et al. PyTorch: An Imperative Style, High-Performance Deep Learning Library. Preprint at 10.48550/arXiv.1912.01703 (2019).

76. Lundberg, S. & Lee, S.-I. A Unified Approach to Interpreting Model Predictions. Preprint at 10.48550/arXiv.1705.07874 (2017).

77. Wolf, F. A., Angerer, P. & Theis, F. J. SCANPY: large-scale single-cell gene expression data analysis. Genome Biol. 19, 15 (2018).

78. Waskom, M. L. seaborn: statistical data visualization. J. Open Source Softw. 6, 3021 (2021).

79. Davidson-Pilon, C. lifelines: survival analysis in Python. J. Open Source Softw. 4, 1317 (2019).

80. Hunter, J. D. Matplotlib: A 2D Graphics Environment. Comput. Sci. Eng. 9, 90–95 (2007).

